# Drug discovery to counteract antinociceptive tolerance with mu-opioid receptor endocytosis

**DOI:** 10.1101/182360

**Authors:** Po-Kuan Chao, Yi-Yu Ke, Hsiao-Fu Chang, Yi-Han Huang, Li-Chin Ou, Jian-Ying Chuang, Yen-Chang Lin, Pin-Tse Lee, Wan-Ting Chang, Shu-Chun Chen, Shau-Hua Ueng, John Tsu-An Hsu, Pao-Luh Tao, Ping-Yee Law, Horace H. Loh, Chuan Shih, Shiu-Hwa Yeh

**Author notes:** Correspondence to: Shiu-Hwa Yeh Institute of Biotechnology and Pharmaceutical Research, National Health Research Institutes, Zhunan 35053, Taiwan. Competing financial interests: The authors have declared that no conflict of interest exists.

## Abstract

Morphine antinociceptive tolerance is highly correlated with its poor ability to promote mu-opioid– receptor (MOR) endocytosis. Our objective was to discover a novel positive allosteric modulator of MOR to enhance morphine-induced MOR endocytosis. We used high-throughput screening to identify several cardiotonic steroids as positive allosteric modulators of morphine-induced MOR endocytosis having high potency and efficacy, independently of Na^+^/K^+^-ATPase inhibition. Convallatoxin was found to enhance morphine-induced MOR endocytosis through an adaptor protein 2/clathrin-dependent mechanism without regulating G protein- or *β*-arrestin-mediated pathways. Both F243 and I292 residues of MOR were essential to the effect of convallatoxin on MOR endocytosis. Co-treatment with chronic morphine and convallatoxin reduced morphine tolerance in animal models of acute thermal pain and chronic inflammatory pain. Acute convallatoxin administration reversed morphine tolerance in morphine-tolerant mice. These findings suggest that cardiotonic steroids are potentially therapeutic for morphine side effects and open a new avenue for the study of MOR trafficking.

## Introduction

Opioids have long been used to treat severe pain (Waldhoer et al., 2004). However, long-term use leads to tolerance, dependence, and addiction (Tao et al., 2010). Opioids primarily activate three GPCRs of the G_i_ subtype: the mu-, delta-, and kappa-opioid receptors (MOR, DOR, and KOR, respectively). Although the mechanisms of opioid-induced analgesia are not well defined, it is now clear that activated opioid receptors are able to utilize both G-protein-dependent and G-protein-independent signaling pathways (Williams et al., 2013). Furthermore, it is generally believed that opioid analgesics exert their pharmacological effects mainly by acting at MOR (Loh et al., 1998).

The general mechanism of opioid tolerance remains controversial (Al-Hasani and Bruchas, 2011); however, unlike other high-efficacy opioids such as [D-Ala^2^, *N*-MePhe^4^, Gly-ol]-enkephalin (DAMGO), etonitazene, etorphine, and fentanyl (Duttaroy and Yoburn, 1995), morphine acted as a poor MOR internalizing agonist (Keith et al., 1996) and developed tolerance through a unique molecular mechanism (Grecksch et al., 2011). Furthermore, morphine tolerance could be limited by enhancing receptor endocytosis (Martini and Whistler, 2007). Previous studies indicated that a mutant MOR with altered recycling (RMOR), which underwent endocytosis after morphine treatment, was associated in vitro with reduced tolerance and with reduced cAMP superactivation, a cellular hallmark of withdrawal (Finn and Whistler, 2001). Compared with WT mice, RMOR knock-in mice showed potentiated morphine antinociception but less tolerance and withdrawal, indicating a beneficial effect of MOR internalization in morphine analgesia (Kim et al., 2008). Furthermore, opioid combinations of morphine with low-dose DAMGO (He et al., 2002) or methadone (He and Whistler, 2005), which are MOR agonists with substantial MOR internalization ability, diminished both morphine tolerance and dependence in rats. However, when using a mixture of agonists, it is difficult to strongly conclude that MOR endocytosis contributes to morphine tolerance/dependence because numerous sets of MOR signaling pathways are modified by each agonist (Alvarez et al., 2002; Blanchet et al., 2003; Enquist et al., 2012; He et al., 2009; Milan-Lobo and Whistler, 2011) according to the so-called agonist-selective theory (Zheng et al., 2010). A previous study indicated that morphine induces MOR endocytosis in mutant L83I (mouse orthologue of human L85I) without altering binding affinity or cAMP signaling. However, cellular tolerance to morphine is reduced (Cooke et al., 2015; Ravindranathan et al., 2009), suggesting an alternative strategy to reduce morphine tolerance by specifically enhancing morphine-induced MOR endocytosis. In this study, we established a high-throughput screening assay to identify signaling-specific positive allosteric modulators (PAMs) to augment morphine-induced MOR endocytosis and validated the ability of the identified molecules to regulate morphine effects in both cell and animal models.

## Results

### Cardiotonic steroids (CTSs) augment morphine-induced MOR endocytosis

To identify PAMs promoting morphine-induced MOR endocytosis, we screened a 480-natural-compound library in the presence of morphine, using a sensitive enzyme complementation assay for MOR endocytosis in human osteosarcoma U2OS cells expressing human MOR (U2OS-MOR). Three CTSs, gitoxigenin, bufalin, and convallatoxin, all capable of inhibiting Na^+^/K^+^-ATPase (Prassas and Diamandis, 2008), were retrieved from the primary screens (Figure 1A, 1B), and their potency and efficacy tested. Concentration-response curves for morphine-induced MOR endocytosis were obtained in the absence or presence at 1 µM of each CTS (Figure 1C). Compared with morphine alone, the maximal response (E_max_) of morphine-induced MOR endocytosis was significantly increased by these three CTSs, but not the half-maximum effective concentration (EC_50_) (Figure 1C). None of the CTSs affected morphine potency (α-factor), but they increased morphine efficacy (β-factor) by 2.8-3.7–fold (Figure 1-Supplementary Table 1). We then determined the potency and efficacy of the CTSs in regulating morphine-induced MOR endocytosis in the presence of the effective concentration of morphine at 10% activation (EC_10_; 0.3 µM), obtained from Figure 1C. The three CTSs enhanced morphine-induced MOR endocytosis in a concentration-dependent manner (Figure 1D), although in the absence of morphine, they themselves induced only slight MOR endocytosis, and only at high concentrations (Figure 1-Supplementary Figure 1). Additional CTSs, including ouabain and digitoxin, also showed significant but weaker effects on enhancing morphine-induced MOR endocytosis (Figure 1-Supplementary Figure 2). We further examined the enhancement of morphine-induced MOR endocytosis by convallatoxin, which is a blood-brain barrier-penetrating CTS (Gozalpour et al., 2014), using live-cell imaging of CHO-K1 cells expressing MOR-CopGFP. Convallatoxin significantly augmented morphine-induced MOR internalization (Figure 1G), whereas morphine or convallatoxin alone had no effect. Additionally, we observed similar results in vivo using immunofluorescent staining for MOR and the plasma-membrane marker wheat germ agglutinin (WGA) in dorsal root ganglion (DRG) neurons obtained from mice co-treated with morphine and convallatoxin (Figure 1H). Moreover, the potentiating effect of convallatoxin was only observed in MOR-expressing cells, indicating an opioid receptor subtype selectivity of CTSs (Figure 1I). In contrast, convallatoxin failed to potentiate the MOR-endocytotic efficacy of methadone, an opioid effective in promoting MOR endocytosis but with a chemical structure distinct from that of morphine (Alvarez et al., 2002), indicating a potential probe dependence of convallatoxin (Figure 1-Supplementary Figure 3). Here, we present the first validation of CTSs as unique, small-molecule enhancers of opioid-induced MOR endocytosis.

**Figure 1.**
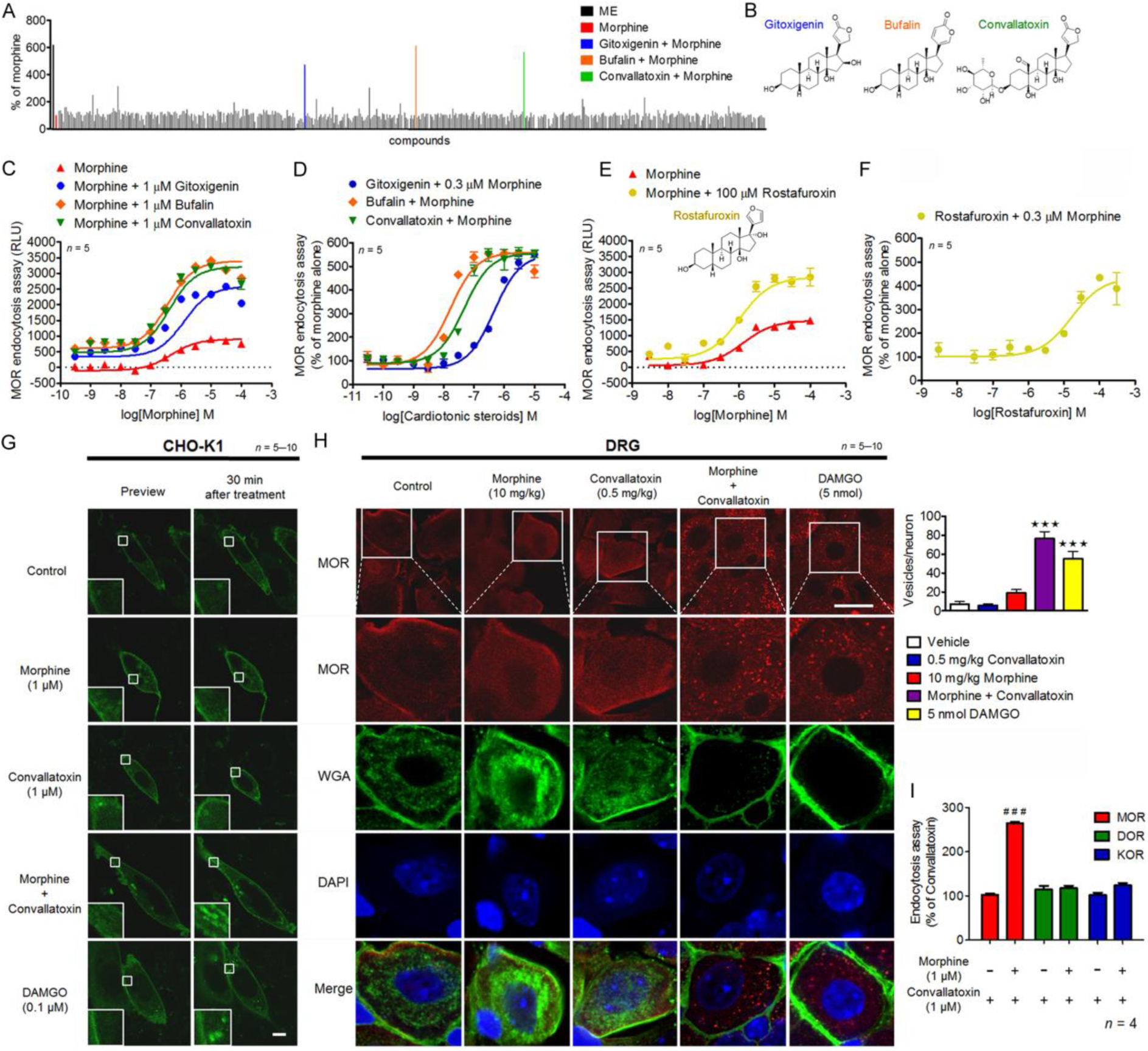
Effect of cardiotonic steroids (CTSs) on opioid-induced mu-opioid receptor (MOR) endocytosis. (**A**) Shown are primary, high-throughput screening results identifying compounds that enhance MOR endocytosis when co-applied with morphine. Heights are percentages of the values for morphine alone at low morphine concentration (0.3 µM; ∼EC10). (**B**) Chemical structures of bufalin, convallatoxin, and gitoxigenin. (**C** and **E**) Concentration-response curves for morphine-induced MOR endocytosis with or without CTSs (**C**) and rostafuroxin (**E**). (**D** and **F**) Concentration-response curves of CTSs (**D**) and rostafuroxin (**F**) in morphine-mediated MOR endocytosis. Shown is varying compound concentration under constant morphine concentration. (**G**) Representative live-cell imaging by real-time confocal microscopy of MOR-CopGFP distribution in CHO-K1 cells before and 30 min after drug treatment. Scale bars, 10 µm. Contents of small boxes are shown at higher magnification in insets. (**H**) Representative immunofluorescence images (left panel) and quantifications (right panel) of the distribution of MOR (red) and wheat-germ agglutinin (WGA, green) in the mouse dorsal root ganglion (DRG) 1 h after drug treatment. The localization of MOR and WGA-labeled plasma membrane is monitored by confocal microscopy. Contents of boxes in top row are shown at higher magnification in second row. DAPI (blue) is a nuclear marker. Scale bar, 20 µm. *F*_4,25_ = 34.75; *p <* 0.001 (1-way ANOVA). (**I**) Receptor subtype-selectivity of CTS. Cells are treated as indicated prior to the receptor-endocytosis assay. *F*_5,18_ = 122; *p <* 0.001 (1-way ANOVA). Data are presented as percentages of the values for convallatoxin alone. Opioid receptor internalization is measured by an enzyme complementation assay in U2OS-MOR, U2OS-DOR, and U2OS-KOR cells. ***, *p* < 0.001 versus vehicle group; ###, *p* < 0.001 versus convallatoxin group (Newman-Keul’s post hoc test). In (**G**–**H**), DAMGO serves as a positive control to induce MOR internalization. All values indicate the mean ± SEM (all figures).

**Figure 2.**
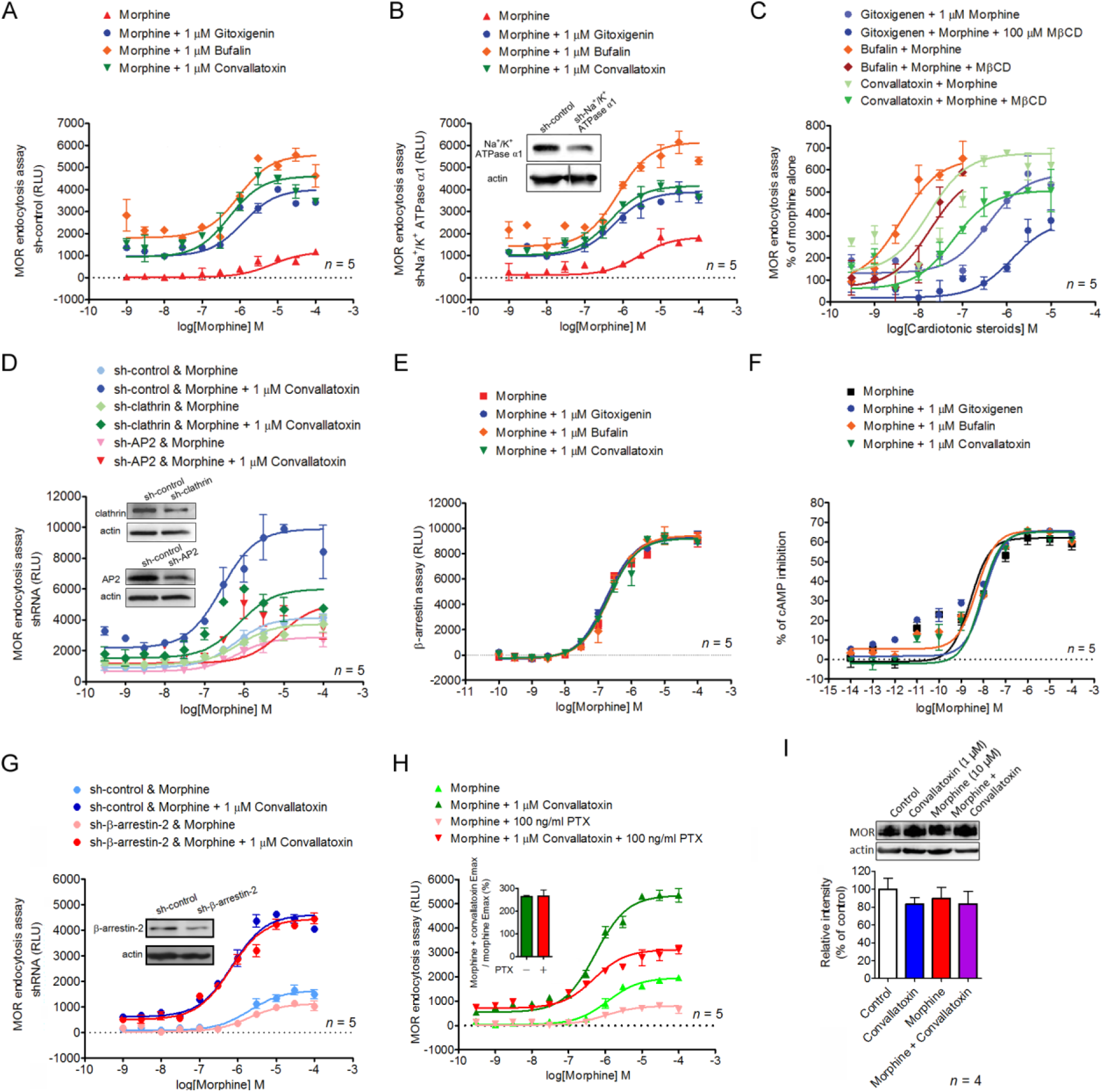
CTSs modulate MOR endocytosis independently of Na^+^/K^+^-ATPase. (**A** and **B**) Role of Na^+^/K^+^-ATPase in the effect of CTSs. U2OS-MOR cells are transiently transfected with *sh-control* (**A**) or *sh-Na*^*+*^*/K*^*+*^*-ATPase α1* (**B,** lower panel), 24 h prior to the MOR internalization assay. (**B**, upper panel) Immunoblot showing *Na*^*+*^*/K*^*+*^*-ATPase α1* expression in knockdown U2OS-MOR cells. (**C**) Concentration-response curves of CTSs in morphine-induced MOR endocytosis in the presence or absence of the endocytosis inhibitor MβCD. Data are percentages of the values for morphine alone (0.3 µM; ∼EC10). (**D**) Silencing of *adaptor protein 2* (*AP2)* or *clathrin* attenuates the effect of convallatoxin on morphine-induced MOR endocytosis. (**D**, lower panel) U2OS-MOR cells are transiently transfected with *sh-control*, *sh-clathrin* or *sh-AP2,* 24 h prior to the MOR internalization assay. (**D**, upper panel) Immunoblots showing *clathrin* or *AP2* expression in *clathrin*- or *AP2-*knockdown U2OS-MOR cells. (**E** and **F**) CTSs fail to modulate morphine-mediated β-arrestin-2 recruitment (**E**) or inhibition of cAMP accumulation (**F**). CHO-K1-MOR cells (**E**) and human embryonic kidney (HEK)-MOR cells (**F**) are treated with various concentrations of morphine in the absence or presence of CTSs. (**G,** lower panel) Silencing of *β-arrestin-2* fails to regulate the effect of convallatoxin on morphine-induced MOR endocytosis. U2OS-MOR cells are transiently transfected with *sh-control* or *sh-β-arrestin-2*, 24 h prior to the MOR internalization assay. (**G**, upper panel) Immunoblot showing *β-arrestin-2* expression in knockdown U2OS-MOR cells. (**H**) Involvement of Gi/o protein in the effect of CTSs on MOR endocytosis. U2OS-MOR cells are pretreated with pertussis toxin 18 h prior to the internalization assay. (**I**) Convallatoxin and/or morphine do not attenuate MOR expression. HEK-MOR cells are treated as indicated for 30 min. Total MOR expression is analyzed by immunoblotting and quantified by densitometry. *F*_3,12_ = 0.45; *p* > 0.05 (1-way ANOVA).

**Figure 3.**
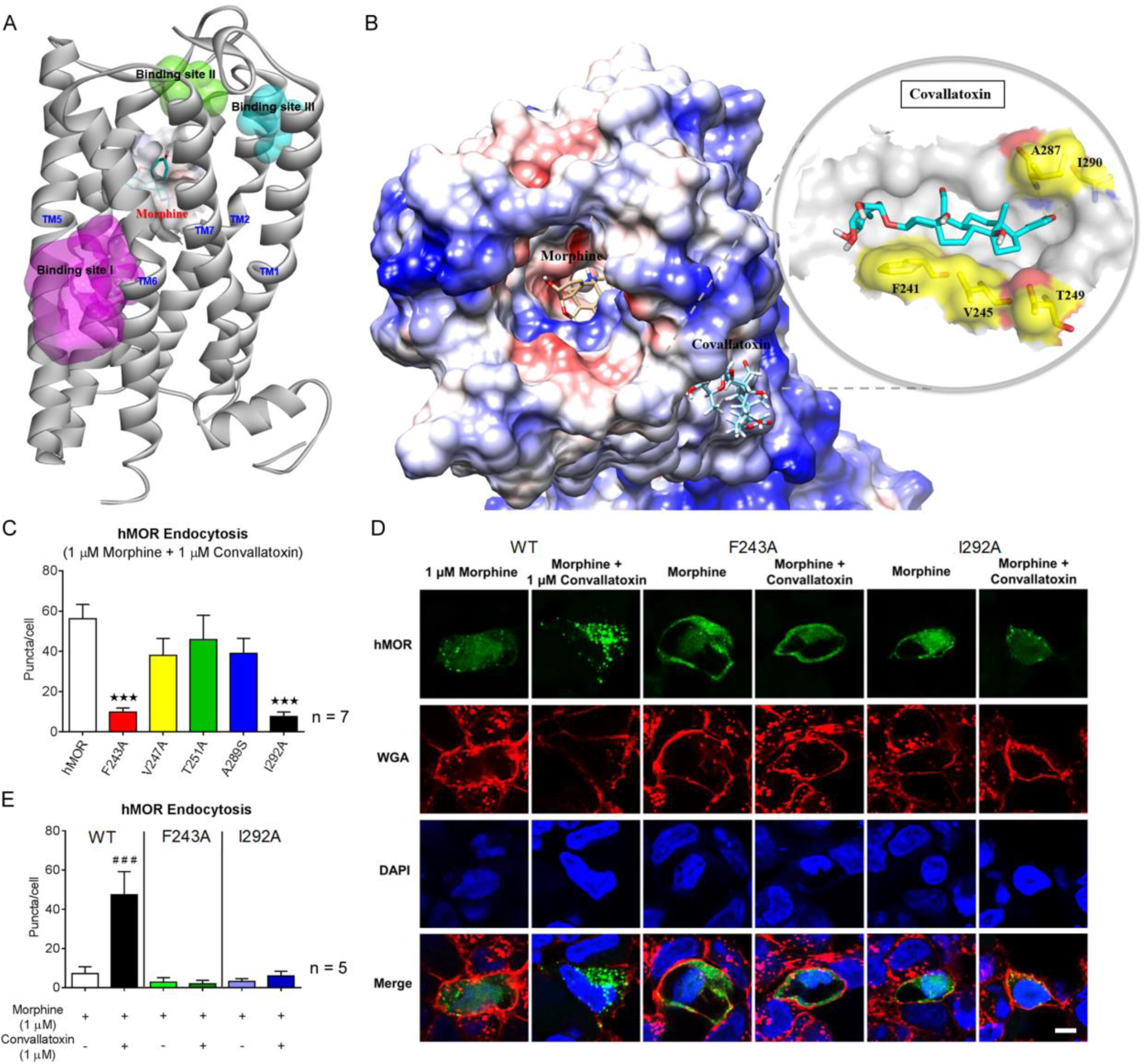
Determining the convallatoxin binding site in MOR. (**A**) The mouse MOR complex with morphine is passed through a 20-ns MD equilibrium calculation, showing release of three binding sites. (**B**) Shown are the binding sites for morphine and convallatoxin and the binding interaction analysis for convallatoxin with mouse MOR. Convallatoxin forms hydrogen bonds with T249 and I290. The hydrophobic effect contributes at F241, V245, A287, and I290. (**C**) Receptor endocytosis analysis of mutants. Cells are treated as indicated (convallatoxin, 1 µM; morphine, 1 µM) prior to live-cell imaging and quantification. *F*_5,24_ = 7.03; *p* < 0.001 (1-way ANOVA). hMOR, human MOR. (**D** and **E**) Representative immunofluorescence images (**D**) and quantification (**E**) of the distribution of CopGFP-tagged MOR and two of its mutants (green) and WGA (red) in mouse dorsal root ganglion (DRG) after drug treatment. *F*_5,24_ = 11.29; *p* < 0.0001 (1-way ANOVA). The localization of MOR and WGA-labeled plasma membrane is monitored by confocal microscopy. DAPI (blue) is a nuclear marker. Scale bar, 20 µm. ***, *p* < 0.001 versus human-MOR group (Newman-Keul’s post hoc test). ###, *p* < 0.001 versus morphine-alone group (Student’s *t*-test).

### CTSs regulate MOR endocytosis through an AP2/clathrin-dependent mechanism

To determine whether the enhancing effect of CTSs on morphine-induced MOR endocytosis is mediated through Na^+^/K^+^-ATPase inhibition, endogenous Na^+^/K^+^-ATPase expression was suppressed by a short, hairpin RNA (shRNA) against the Na^+^/K^+^-ATPase α1 subunit (*shNa*^*+*^*/K*^*+*^*ATPase α1*) in U2OS-MOR cells. Each CTS continued to enhance morphine efficacy (β-factor) by 2.3-5.2-fold in cells transfected either with the universal negative control shRNA (sh-control) or with *shNa*^*+*^*/K*^*+*^*ATPase α1* (Figure 2A, 2B). Furthermore, treatment of U2OS-MOR cells with rostafuroxin, a CTS without Na^+^/K^+^-ATPase inhibitory activity (Ferrari et al., 2006), also enhanced morphine efficacy (β-factor) by 2-fold (Figure 1E, 1F). These results indicate that CTSs potentiate morphine-induced MOR endocytosis through a Na^+^/K^+^-ATPase-independent mechanism. However, pretreatment of cells with the endocytosis inhibitor methyl-β-cyclodextrin (MβCD) (Rodal et al., 1999), or the silencing of two genes generally involved in receptor endocytosis, *adaptor protein 2* (*AP2*) and *clathrin*, attenuated the enhancing effects of CTSs (Figure 2C, 2D), indicated that the machinery of endocytosis is critical for the CTS’s enhancing effect.

We evaluated the ability of CTSs to alter three signaling pathways: MOR-mediated G protein-independent signaling (*β*-arrestin recruitment), G protein-dependent signaling (adenylyl cyclase inhibition), and G protein-coupled inwardly rectifying potassium (GIRK) channel activation (another G protein-dependent signaling pathway known to contribute to MOR-mediated analgesia) (Marker et al., 2004; Nockemann et al., 2013). The CTSs failed to modulate morphine-induced *β*-arrestin-2 recruitment to MOR as determined by the PathHunter enzyme complementation assay in CHO-K1 cells expressing human MOR (CHO-K1-MOR) (Figure 2E, Figure 1-Supplementary Table 1). They also failed to modulate morphine-induced inhibition of cAMP production as determined by a cAMP assay in human embryonic kidney 293 (HEK-293) cells constitutively expressing human MOR (HEKMOR) (Figure 2F, Figure 1-Supplementary Table 1). They likewise failed to modulate morphine-induced, MOR-dependent membrane potential hyperpolarization as determined by a membrane potential assay in mouse pituitary AtT-20 cells transiently expressing human MOR (Figure 2-Supplementary Figures 1, 2A). Furthermore, the potentiation effect of CTSs on MOR endocytosis was retained both in *β*-arrestin-2 knockdown cells (Figure 2G), and in cells pretreated with the receptor-Gi/o protein uncoupler, pertussis toxin (Figure 2H). Expression of MOR protein was not affected either by convallatoxin, morphine, or their combination (Figure 2I). Thus, CTSs did not alter morphine-induced β-arrestin-2 recruitment, adenylyl cyclase inhibition, or GIRK-channel activation. Furthermore, the effect of CTSs on MOR endocytosis was retained even after these two pathways were inhibited. However, CTSs did enhance morphine-induced MOR endocytosis in an AP2/clathrin-dependent manner without altering protein expression.

### Identification of the convallatoxin binding site on MOR

To analyze the potential binding site of CTSs on MOR, the co-crystal structure of opioid agonist BU72 and mouse MOR was applied to computational docking studies (Huang et al., 2015b; Sounier et al., 2015). We first predicted the morphine binding conformation. After 20 ns of equilibrium molecular dynamics, morphine revealed a binding interaction with MOR different from that existing before equilibration (Figure 3-Supplementary Figure 1). This resulted in a conformational change at MOR that exposed further binding sites (binding sites I, II, and III) surrounding the MOR transmembrane (TM) helices 1, 2, 5, 6, and 7 (TM1, TM2, TM5, TM6, and TM7; Figure 3A). Convallatoxin was then docked to these binding sites to identify the most favorable binding residues. According to the Analyze Ligand Poses protocol of the Discovery Studio 2016 application, convallatoxin forms hydrogen bonds with T249 and I290, and generates hydrophobic interactions with F241, V245, A287, and I290, all located in TM5 and TM6 of mouse MOR (Figure 3-Supplementary Table 1, Figure 3-Supplementary Figure 2A). Moreover, these residues are all located in binding site I, and a possible binding mode for convallatoxin based on these results is illustrated in Figure 3B. Sequence conservation analysis demonstrates good conservation between mouse and human for all key binding-site residues (Figure 3-Supplementary Figure 2B).

To test the modelled convallatoxin site identified in the structure, F243, V247, T251, or I292 (human orthologues of F241, V245, T249, and I290) were mutated to alanine, and A289 (human orthologue of A287) was mutated to serine, to determine their roles in morphine-mediated MOR endocytosis. The mutations were introduced into the full-length, WT receptor with a C-terminal–CopGFP tag. The presence of the CopGFP tag had no impact on morphine activation of the receptor (data not shown). G-protein–dependent signaling (GIRK activation) is retained in all five mutant receptors after morphine treatment (Figure 2-Supplementary Figure 2B–2F). However, mutation of F243 and I292 to alanine significantly reduced the effect of convallatoxin, whereas convallatoxin continued to potentiate morphine-mediated MOR endocytosis in the mutations of V247 and T251 to alanine, and of A289 to serine, both in CHO-K1 cells (Figure 3C) and DRG neurons (Figure 3D–3E). Thus, these results support a role for F243 and I292 residues in the functions of MOR endocytosis, and are consistent with the modelled binding site for convallatoxin.

### Convallatoxin treatment diminishes morphine tolerance in an AP2/clathrin-dependent manner in mice

To assess the effect of convallatoxin on morphine-produced ant nociception, tail-flick tests after acute and chronic treatments were performed as shown in Figure 4A–4D. Acute morphine and morphine + convallatoxin displayed a similar potency (ED_50_: morphine, 2.6 ± 0.3 mg/kg; morphine + convallatoxin, 2.8 ± 0.7 mg/kg) and magnitude of antinociception (Figure 4A, 4C). However, chronic morphine + convallatoxin resulted in greater potency (ED_50_: morphine, 10.6 ± 1.2 mg/kg; morphine + convallatoxin, 6.4 ± 1.2 mg/kg) and magnitude (Figure 4D) of antinociception, and greater MOR endocytosis (Figure 4G) relative to chronic morphine alone. These results indicate that chronic treatment with convallatoxin reduces morphine antinociceptive tolerance. The experiments described in Figure 4A–4D were then conducted in MOR-KO mice. No difference in the response to acute treatment (Figure-Supplementary Figure 1A) and chronic treatment (Figure 4-Supplementary Figure 1B) was found between study groups, suggesting that the effects of convallatoxin on morphine antinociception are mediated through MOR signaling.

Tolerance may be unavoidable after long-term use of morphine (Labianca et al., 2012), and a strategy to reverse these characteristics should be of benefit for chronic pain management (Becker, 2010). We therefore investigated the effects of acute CTS administration on morphine-tolerant mice receiving chronic morphine injection twice daily for 8 d. On the test day, mice were challenged with morphine, either alone or in combination with convallatoxin, and tested for antinociception with the tail-flick model (Figure 4E). Morphine-tolerant mice that received acute morphine with convallatoxin showed significantly greater antinociception (Figure 4F) than mice that received morphine alone. Furthermore, the dosage of convallatoxin we applied did not change the basal locomotor activity in the open field test (Figure 4-Supplementary Figure 2). Thus, these results indicated that acute CTS treatment was able to reverse morphine antinociceptive tolerance.

**Figure 4.**
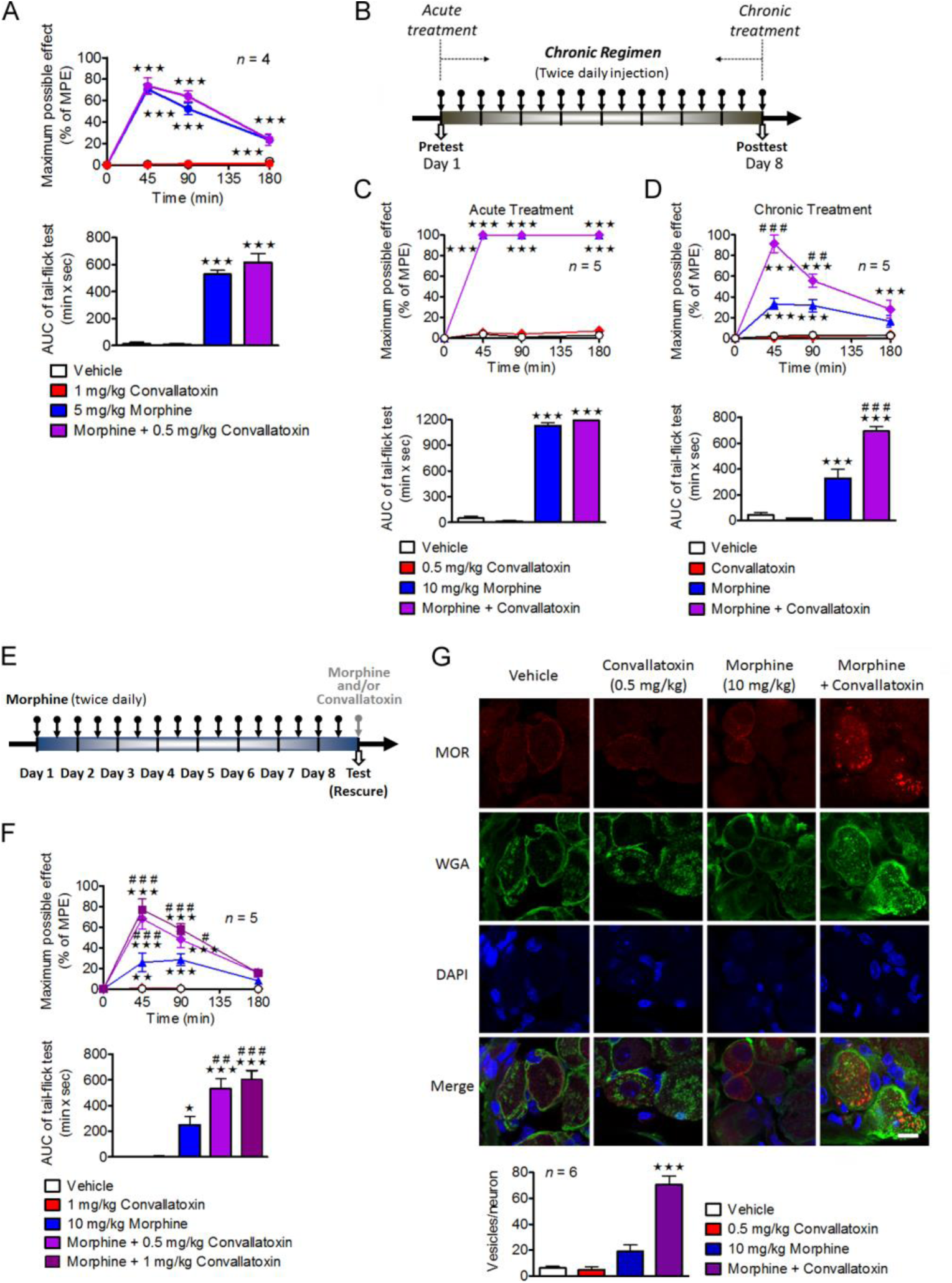
Convallatoxin enhances MOR endocytosis in vivo and diminishes tolerance to morphine antinociception. (**A**) Effect of convallatoxin on morphine antinociception. (upper panel) Treatment *F*_3,9_ = 105.9, time *F*_3,9_ = 82.97, interaction *F*_9,27_ = 37.87; *p <* 0.001 (2-way ANOVA). Quantitative results from time-response curves are presented as AUCs. (lower panel) *F*_3,12_ = 82.9; *p <*0.001 (1-way ANOVA). (**B**) Experiment flowchart for investigating the effect of convallatoxin on the development of morphine tolerance. (**C** and **D**) Acute (**C**) and chronic (**D**) antinociceptive effects of each treatment in mice measured using the tail-flick test. (**C**, upper panel) Treatment *F*_3,16_ = 1938, time *F*_3,48_ = 3828, interaction *F*_9,48_ = 1089; (**D**, upper panel) Treatment *F*_3,16_ = 80.31, time *F*_3,48_ = 41.5, interaction *F*_9,48_ = 18.41; all *p <* 0.001 (2-way ANOVA). Quantitative results from time-response curves are presented as AUCs. (**C**, lower panel) *F*_3,16_ = 1068; (**D**, lower panel) *F*_3,16_ = 62.45; all *p <* 0.001 (1-way ANOVA). (**E**) Experiment flowchart for investigating the effect of acutely administered convallatoxin in morphine-tolerant mice. (**F**) The acute antinociceptive effects of each treatment in morphine-tolerant mice. Treatment *F*_4,20_ = 30.24, time *F*_3,60_ = 81.9, interaction *F*_12,60_ = 18.07; all *p <* 0.001 (2-way ANOVA; upper panel). Quantitative results from time-response curves, presented as AUCs. *F*_4,20_ = 26.54; *p <* 0.001 (1-way ANOVA; lower panel). (**G**) Representative immunofluorescence images (upper panel) and quantification (lower panel) of MOR (red) and WGA (green) distribution in the mouse DRG after chronic drug treatment. DAPI (blue) is a nuclear marker. Scale bar, 20 µm. *F*_3,20_ = 52.02; *p <* 0.001 (1-way ANOVA). Data in (**A**, upper panel), (**C**, upper panel), (**D**, upper panel), (**F**, upper panel): **, *p* < 0.01; ***, *p* < 0.001 versus vehicle group; #, *p* < 0.05; ##, *p* < 0.01; ###, *p* < 0.001 versus morphine-alone group (Bonferroni’s post hoc test). Data in (**A**, lower panel), (**C**, lower panel), (**D**, lower panel), (**F**, lower panel), (**G**, lower panel): *, *p* < 0.05; ***, *p* < 0.001 versus vehicle group; ##, *p* < 0.01; ###, *p* < 0.001 versus morphine-alone group (Newman-Keul’s post hoc test). MPE, maximum possible effect.

Because we found that convallatoxin enhances MOR endocytosis through an AP2/clathrin pathway, we then examined the role of AP2/clathrin on the effect of convallatoxin in morphine antinociception. Intrathecal electroporation of shRNA silenced either clathrin or AP2 in DRG neurons (Figure 5-Supplementary Figure 1) without influencing the basal nociceptive sensitivity of the mice. However, the enhancement effect of chronic convallatoxin on morphine antinociception (Figure 5A–5B) and MOR endocytosis (Figure 5C) were abolished in the spinal AP2 or clathrin knockdown mice, suggesting that the AP2/clathrin adaptor complex mediates the chronic effect of convallatoxin in vivo.

**Figure 5.**
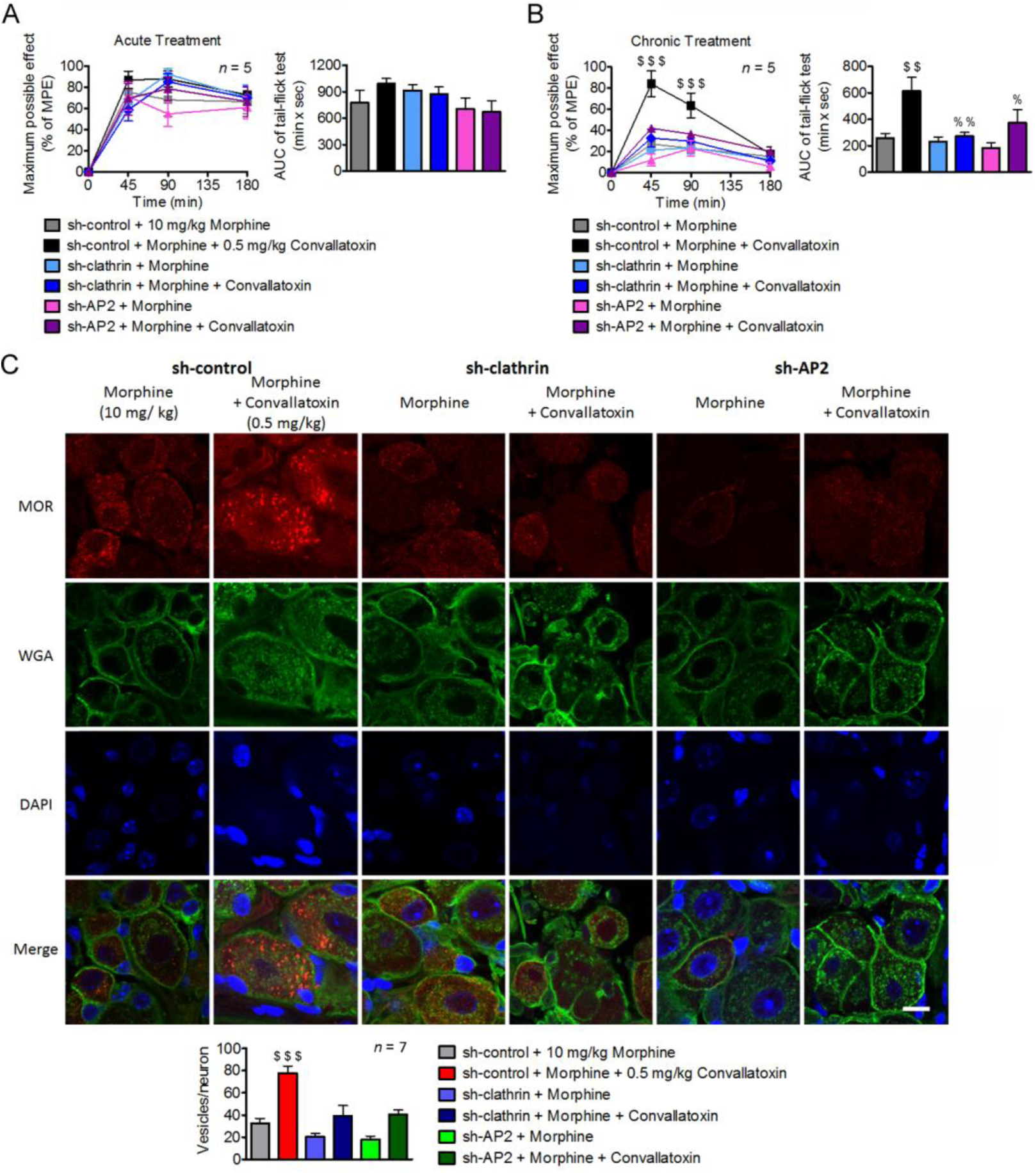
Down-regulation of clathrin and AP2 attenuated the effect of convallatoxin in morphine antinociceptive tolerance. (**A** and **B**) The *sh-control*, *sh-clathrin*, or *sh-AP2* plasmids are delivered into the spinal cord of WT mice using direct in vivo electroporation. Seven days after surgery, the antinociceptive effects after acute (**A**) and chronic (**B**) treatment with drugs are measured using the tail-flick test. (**A**, left panel) Treatment F5,24 = 0.47, time F3,72 = 168.2, interaction F15,72 = 1.44; (**B**, left panel) Treatment F5,24 = 6.9, time F3,72 = 72.56, interaction F15,72 = 5.81; all *p* < 0.001 (2-way ANOVA). Quantitative results from the left panel of (**A**) and (**B**) are presented as AUCs. (**A**, right panel) F5,24 = 1.43; *p* > 0.05; (**B**, right panel) F5,24 = 5.64; *p* < 0.01 (1-way ANOVA). (**C**) Representative immunofluorescence images (upper panel) and quantification (lower panel) of MOR (red), and WGA (green) for each treatment in mouse DRG neurons. DAPI (blue) is a nuclear marker. Scale bars, 20 µm. *F*_5,36_ = 14.47; *p <* 0.001 (1-way ANOVA). Data in (**B**, left panel): $$$, *p* < 0.001 versus *sh-control* + morphine group (Bonferroni’s post hoc test). Data in (**B**, right panel), (**C**, lower panel): $$, *p* < 0.01; $$$, *p* < 0.001 versus sh-control + morphine group; %, *p* < 0.05; %%, *p* < 0.01 versus sh-control + morphine + convallatoxin group (Newman-Keul’s post hoc test). MPE, maximum possible effect.

### Convallatoxin treatment reduces morphine tolerance in mice with chronic inflammatory pain

Opioids are used to manage chronic osteoarticular pain, but repeated administration results in the development of antinociceptive tolerance (Fernández-Dueñas et al., 2007). We thus investigated whether a CTS has beneficial effects on morphine antinociception in the complete Freund’s adjuvant (CFA)-induced mouse model of rheumatoid arthritis (Nagakura et al., 2003), and mechanical allodynia was measured by the Von Frey test to examine the antinociceptive effects of each treatment on post-inoculation days (PIDs) 14, 16, and 18 (Figure 6A). The threshold of mechanical allodynia in CFA-treated mice decreased significantly during PIDs 14 to 18 compared with saline-treated mice and reached a plateau. In CFA-treated mice, acute treatment with morphine + convallatoxin, but not with morphine alone, on PID 14 diminished mechanical allodynia, and the paw withdrawal threshold was similar to that in the non-CFA-treated groups (Figure 6A). Furthermore, the morphine + convallatoxin group showed reduced antinociceptive tolerance relative to the morphine group. After daily treatment with morphine or morphine + convallatoxin for 5 d continuously (PID 18), the threshold of mechanical allodynia of these two experimental groups was reduced by approximately 92% (Figure 6B) and 67% (Figure 6C), respectively, in CFA-treated mice, whereas convallatoxin itself did not produce any antinociceptive effect. Moreover, the silencing of *clathrin* and *AP2* in DRG neurons of B6 mice significantly decreased the effect of convallatoxin on both acute and chronic morphine antinociception (Figure 6D), as well as on MOR endocytosis (Figure 6E). This further supports the conclusion that CTSs enhance morphine efficacy and attenuate the development of antinociceptive tolerance through an AP2/clathrin-dependent pathway in chronic inflammatory pain.

## Discussion

CTSs are used to treat cardiac failure and atrial fibrillation through inhibition of Na^+^/K^+^-ATPase (Wehrens and Marks, 2004). In addition to acting as Na^+^/K^+^-ATPase inhibitors, CTSs may regulate other proteins, potentially enabling them to serve as therapeutic agents. Bufalin selectively reduces the protein levels and intrinsic transcriptional activity of steroid receptor coactivators (SRC)-1 and SRC-3, and has potential as a broad-spectrum cancer inhibitor (Wang et al., 2014). Bufalin has also been shown to inhibit interferon-β expression and tumor necrosis factor signaling, and has thus been proposed as a treatment for inflammatory and autoimmune diseases, respectively (Ye et al., 2011).

**Figure 6.**
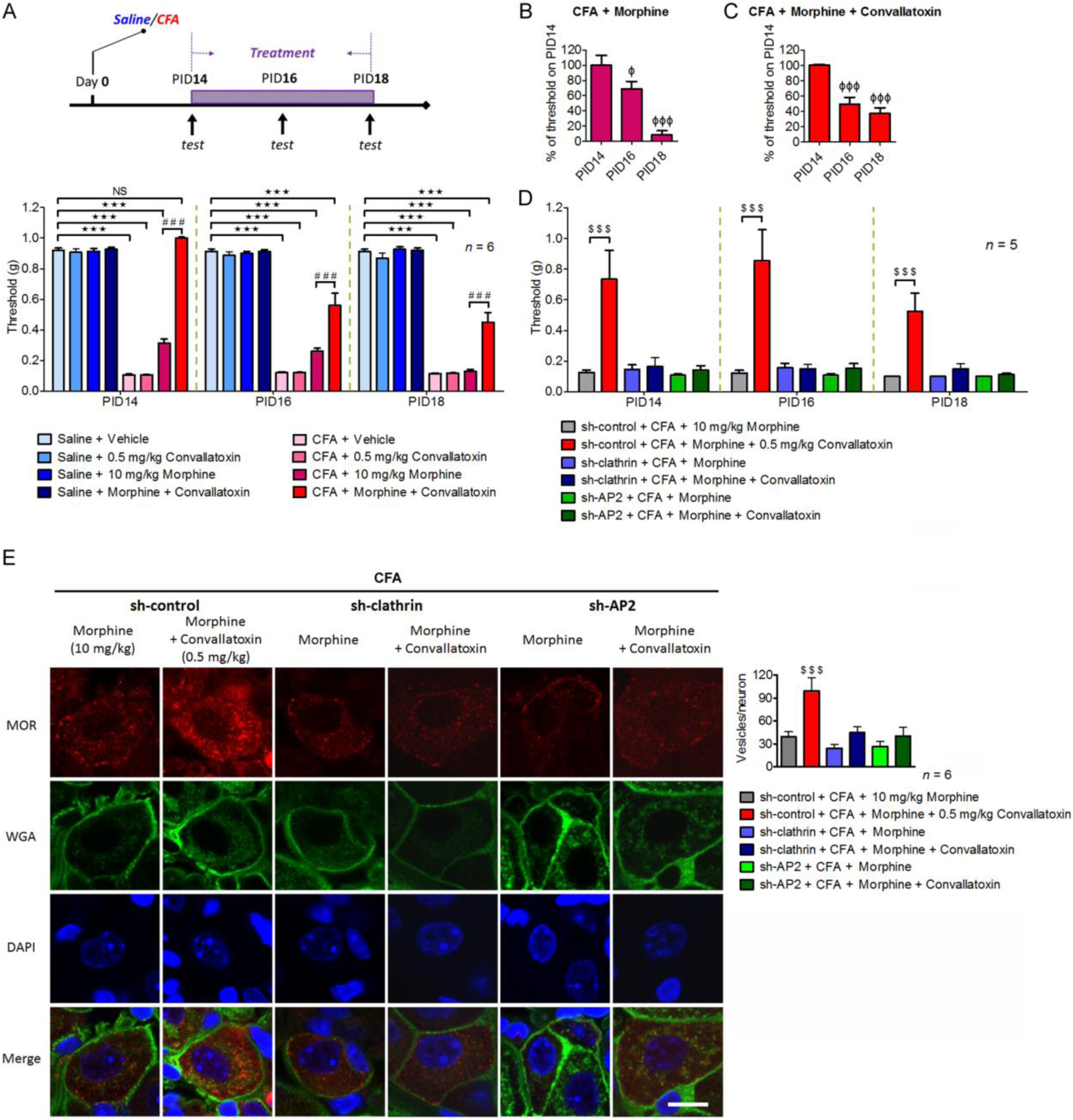
Convallatoxin alters morphine-induced alleviation of mechanical allodynia through clathrin and AP2 in the CFA-induced arthritic mouse model. (**A**) Experimental flowchart for effects of convallatoxin on allodynia. Mice receive an intraplantar injection of saline or CFA to induce local inflammation. Allodynia, expressed in g, is evaluated in CFA- and saline-treated mice 45 min after the last drug injection on post-inoculation days (PIDs) 14, 16, and 18. Treatment *F*_7,40_ = 384.2, time *F*_2,80_ = 40.7, interaction *F*_14,80_ = 23.48; all *p <* 0.001 (2-way ANOVA). (**B** and **C**) Quantitative results from (**A**) are presented as percentages of the threshold on PID 14 for CFA + morphine (**B**) or CFA + morphine + convallatoxin (**C**). (**B**) *F*_2,15_ = 22.32; (**C**) *F*_2,15_ = 24.7; all *p <* 0.001 (1-way ANOVA). (**D**) Silencing *clathrin* or *AP2* attenuates the effect of convallatoxin on morphine antinociception. The mouse spinal cord was electroporated with *sh-control*, *sh-clathrin*, or *sh-AP2* on PID 7. Treatment *F*_5,24_ = 58.27, time *F*_2,48_ = 1.32, interaction *F*_10,48_ = 0.58; all *p <* 0.001 (2-way ANOVA). (**E**) Representative immunofluorescence images (left panel) and quantification (right panel) of MOR (red) and WGA (green) for each treatment in mouse DRG neurons. DAPI (blue) is the nuclear marker. Scale bar, 20 µm. *F*_5,30_ = 7.36; *p <* 0.001 (1-way ANOVA). Data in (**A**, lower panel) and (**D**): ***, *p* < 0.001 versus saline + vehicle group; ###, *p* < 0.001 versus CFA + morphine group; $$$, *p* < 0.001 versus *sh-control* + CFA + morphine group (Bonferroni’s post hoc test). Data in **B**, **C** and **E**, right panel: Φ, *p* < 0.05; ΦΦΦ, *p <* 0.001 versus threshold of each group on PID 14; $$$, *p* < 0.001 versus *sh-control* + CFA + morphine group (Newman-Keul’s post hoc test).

In this study, we first identified CTSs as novel PAMs of morphine-induced MOR endocytosis (Figure 1) independently of adenylyl cyclase inhibition, β-arrestin-2 recruitment (Figure 2E–2H), and Na^+^/K^+^-ATPase inhibition (Figure 2A, 2B), which is different from previously known PAMs of MOR (BMS-986121 and BMS-986122), which show an absence of signaling bias (Burford et al., 2013). Furthermore, convallatoxin prevented the development or expression of morphine tolerance in a MOR-dependent manner in vivo (Figures 4, 5), and enhanced morphine antinociception in mice with CFA-induced hind-paw allodynia (Figure 6).

Searching for allosteric modulators of GPCRs is a prominent topic for next-generation drug discovery. MOR-selective PAMs for the cAMP and β-arrestin pathways have been identified, and previous studies have indicated that morphine can selectively induce endocytosis in the mutant L83I (rat orthologue of human L85I) MOR without changing receptor binding affinity or the cAMP pathway, suggesting the existence of PAMs for MOR endocytosis. In the present study, CTSs bound to the F243 and I292 residues located in human MOR TM5 and TM6, which are far from L83, to potentiate morphine-mediated MOR endocytosis, potentially indicating a new allosteric site for MOR endocytosis. According to docking and molecular-dynamical analysis using mouse MOR, the I290 residue is suggested to form a hydrogen bond with the carbonyl group of the furanone of convallatoxin, and F241 is supposed to engage in pi-alkyl hydrophobic interactions with the methyl substituent on the tetrahydropyrantriol moiety and with the methylene group on the second carbon of convallatoxin (Figure 6-Supplementary Figure 1). These findings warrant further investigation. On the other hand, one could imagine that PAMs of MOR might favor signaling pathways related to antinociception and disfavor signaling pathways that mediate unwanted side effects. Such abilities have been observed in allosteric modulators of the muscarinic system (Davis et al., 2009). In the present study, CTSs selectively potentiate morphine-mediated MOR endocytosis without altering the cAMP or β-arrestin pathways, suggesting a new class of PAMs distinguished by signaling bias. Another compelling characteristic of CTSs is their ability to enhance the efficacy (β factor) rather than the potency (α factor) of morphine. In the future, drug development programs will benefit from this behavior because it eliminates the need to change the dosage of morphine when combining it with CTSs.

Upon agonist-induced activation of MOR, G-protein coupling (Zaki et al., 2000) and β-arrestin-2 recruitment (Whistler and von Zastrow, 1998) are believed to be essential for promoting receptor internalization through a linkage of GPCRs to proteins of the endocytotic machinery, including clathrin (Goodman et al., 1996) and AP2 (Laporte et al., 1999). However, morphine-induced G-protein coupling and β-arrestin-2 recruitment were not altered by CTSs in the present study (Figure 2E, 2F), and blockade of these pathways failed to attenuate the effects of CTSs on morphine-mediated MOR endocytosis (Figure 2G, 2H). In fact, recent research has shown that β-arrestin-2 recruitment is not necessary for internalization of GPCRs. Several neuronal GPCRs, including the metabotropic glutamate receptor 1 (Dhami et al., 2004), the serotonin 5-HT2A receptor (Gray et al., 2003), and the M2 muscarinic cholinergic receptor (van Koppen and Kaiser, 2003), show β-arrestin-independent receptor internalization. Since clathrin and AP2 are involved with the internalization of numerous proteins, and since CTSs potentiated MOR endocytosis rather than that of DOR or KOR (Figure 1I), a MOR-specific mechanism for CTSs seems possible (Ritter and Hall, 2009; Soohoo and Puthenveedu, 2013). Furthermore, CTSs enhanced MOR endocytosis through a β-arrestin-2-independent and AP2/clathrin-dependent mechanism, which warrants further investigation.

We investigated the effects of CTSs on acute and chronic morphine in animal models of acute thermal and chronic inflammatory pain. Convallatoxin significantly prevented the development of chronic, morphine-induced tolerance. Additionally, acute convallatoxin administration reversed morphine tolerance in morphine-tolerant mice (Figure 4). In CFA-induced chronic inflammatory pain, morphine only partially relieved the mechanical allodynia (Figure 6A). These results agree with previous studies showing that MOR agonists are highly effective in acute pain, but less so in chronic inflammatory and neuropathic pain (Fernández-Dueñas et al., 2007; Mankovsky et al., 2012; Rowbotham et al., 2003). Interestingly, the morphine-induced decrease in mechanical allodynia was enhanced by convallatoxin through AP2/clathrin-dependent pathways (Figure 6), which introduces the possibility that morphine’s poor ability to induce MOR endocytosis underlies its lack of efficacy in treating chronic inflammatory pain (Kim et al., 2008). It is worth noting that convallatoxin itself exerted no antinociception (Figures 4A, 4C, 4D, 4F, and 6A), and that the effects of convallatoxin on morphine-produced antinociception were attenuated in mice with spinal *AP2* or *clathrin* knockdown (Figures 5, 6D, 6E). In contrast, two other CTSs, ouabain and digitoxin, which have greater potency as Na^+^/K^+^-ATPase inhibitors than does convallatoxin (Lamers et al., 1985), acted as weak enhancers of morphine-induced MOR endocytosis relative to other glycosides (Figure 1-Supplementary Figure 2). However, they have been shown to produce antinociception themselves or to antagonize morphine-induced antinociception (Gonzalez et al., 2012; Masocha et al., 2003; Zeng et al., 1999). It is unlikely that the Na^+^/K^+^-ATPase inhibition ability of CTSs is associated with ligand-induced MOR endocytosis because knockdown of Na^+^/K^+^-ATPase failed to attenuate the effects of CTSs on morphine-induced MOR endocytosis (Figure 2A, 2B). Furthermore, another CTS, rostafuroxin, which does not inhibit Na^+^/K^+^-ATPase, also enhanced morphine-induced MOR endocytosis (Figure 1E, 1F). We thus suggest that other than Na^+^/K^+^-ATPase inhibition, CTSs potentially have a structure-activity relationship related to ligand-induced MOR endocytosis that correlates with MOR-mediated analgesia. Therefore, it should be beneficial to modify the chemical structures of CTSs to increase ligand-induced MOR analgesia by enhancing ligand-induced MOR endocytosis, while diminishing cardiotoxic effects by reducing Na^+^/K^+^-ATPase inhibition. A similar strategy has been used for structural modification of digoxin to reduce cytotoxicity while maintaining inhibitory effects on T-cell differentiation (Huh et al., 2011).

In conclusion, our findings demonstrate that cardiotonic steroids act as novel positive allosteric modulators specifically enhancing morphine-induced MOR endocytosis, which enhances opioid antinociception and decrease morphine tolerance (Figure 7). These studies provide proof-of-concept for the development of signaling-specific opioid allosteric modulators, as well as new insights into clinical opioid-managed pain. The newly discovered regulatory mechanism of ligand-induced MOR endocytosis opens a new avenue in the study of the trafficking of MOR and of other GPCRs.

**Figure 7.**
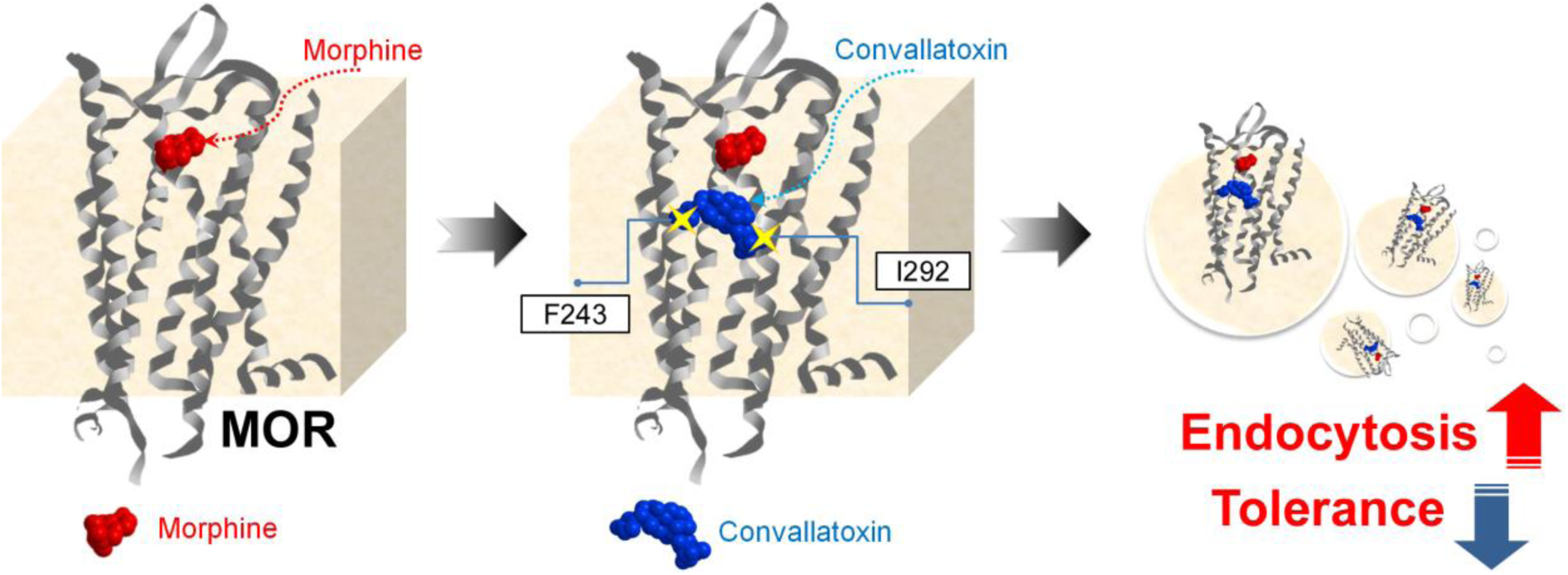
CTSs are the first-discovered, small-molecule, positive allosteric modulators for morphine-induced MOR endocytosis. Morphine binds to MOR and elicits antinociceptive effects. However, its poor ability to induce MOR endocytosis results in morphine tolerance after long-term use. CTSs bind to the F243 and I292 residues, induce a conformational change in MOR, and thus promote better induction of receptor endocytosis by morphine, accompanied by better antinociception and reduced morphine tolerance.

## Materials and methods

### Animals

Male, eight- to ten-week-old WT C57BL/6 (B6) mice (25–30 g) and MOR-KO mice (Chen et al., 2010) (provided by Dr. Pao-Luh Tao, National Health Research Institutes, Taiwan) were kept in a temperature-controlled animal room with a 12:12 h light-dark cycle.

### shRNA transfection and site-directed mutagenesis

The pLKO.1 vectors encoding shRNA directed against the Na^+^/K^+^-ATPase α1 subunit (Clone ID: TRCN0000332624), β-arrestin-2 (Clone ID: TRCN0000165387), clathrin heavy polypeptide (Clone ID: TRCN0000113160), AP2 α1 subunit (Clone ID: TRCN0000065112), and the universal negative control (Clone ID: TRCN0000231759) were from the National RNAi Core Facility (Institute of Molecular Biology/Genomic Research Center, Academia Sinica, Taiwan). Cells were transfected with the shRNA using TurboFect transfection reagent (Thermo Scientific). Twenty-four hours after transfection, cells were harvested for the internalization assay, and protein levels were determined using immunoblotting. For site-directed mutagenesis, the human wild-type MOR gene was located in a pCDH-EF1-MCS-T2A-copGFP plasmid (System Biosciences) with C-terminal CopGFP tag, and MOR mutants were generated using the QuikChange Lightning Site-Directed Mutagenesis Kit (Agilent Technologies) and appropriate primers (Figure 3-Supplementary Table 2). The detail experimental procedures were following manufacturer’s instructions.

### Live cell imaging

CHO-K1 cells (ATCC) were grown in F12 medium containing 10% FBS, 100 units/mL penicillin, and 100 μg/mL streptomycin in T-175 tissue culture flasks and harvested with trypsin/EDTA solution. Cells were transiently transfected with the wild-type or mutant human *MOR-copGFP* plasmids using the NEPA21 electroporator gene transfection system (Nepa Gene) and were subsequently seeded into a glass-bottomed culture dish (MatTek Corporation). The pulse conditions for electroporation were as follows: 125 V, pulse length 7.5 ms, inter-pulse interval 50 ms, decay rate 10%, and polarity plus. The transfer pulse conditions were as follows: 20 V, pulse length 50 ms, pulse interval 50 ms, decay rate 40%, and polarity plus and minus. After 24 h, cells were serum-starved for 3 h and then MOR trafficking was measured. Cells were treated with vehicle, morphine (1 µM; Food and Drug Administration, Ministry of Health and Welfare, Taipei, Taiwan), convallatoxin (1 µM; Sigma), morphine (1 µM) + convallatoxin (1 µM), or DAMGO (0.1 µM; Tocris Biosciences) for 30 min, and images were captured with a laser confocal microscope (TCS SP5II, Leica) using the same gain and exposure time for each group. Images were then superimposed to determine the localization of the MORs.

### Immunoblotting

Cells were lysed in homogenization buffer (1% Triton X-100, 50 mM Tris-HCl, pH 7.5, 0.3 M sucrose) containing protease and phosphatase inhibitor cocktails (Roche). Equal amounts of the samples (20–50 µg) were resolved using 8.5% or 10% SDS-PAGE and transferred to Immobilon-P membranes (Millipore). Blots were incubated with rabbit monoclonal anti-MOR antibody (1:1000; Cat. NBP1-96656; Novus Biologicals), mouse monoclonal anti-Na^+^/K^+^-ATPase α1 subunit antibody (1:1000; Cat. ab7671; Abcam), rabbit monoclonal anti-AP2 α antibody (1:1000; Cat. GTX62588; GeneTex), mouse monoclonal anti-clathrin heavy chain antibody (1:1000; Cat. GTX22731; GeneTex), or mouse monoclonal anti-β-actin antibody (1:5000; Cat. A5316; Sigma) for 16 h. Following incubation, blots were washed and incubated in horseradish peroxidase-conjugated secondary antibody (1:1000; Cat. NA931V, NA934V; GE Healthcare) for 1 h. Signals were detected using a chemiluminescence enhancement kit (Millipore), and the density of the immunoblots was determined using ImageJ (NIH).

### Immunostaining

B6 mice were first injected subcutaneously (s.c.) with vehicle, morphine (10 mg/kg), convallatoxin (0.5 mg/kg), morphine (10 mg/kg) + convallatoxin (0.5 mg/kg), or intrathecally (i.t.) with DAMGO (5 nmol), for 60 min, and then anesthetized by isoflurane, sacrificed, and perfused transcardially with 1 × PBS followed by 4% paraformaldehyde. The DRG neurons were removed and postfixed for 12 h in 4% paraformaldehyde and then cryoprotected in 20% glycerol for 6 h at 4°C. The DRG sections were glued to the platform of a Vibroslice tissue slicer. Transverse sections of 20-μm thickness were cut and the appropriate slices from each group were placed on the same microscope slide and processed identically during a standard immunofluorescence staining procedure. Double-labeling immunohistochemistry was performed as previously described (Lee et al., 2014). Sections were first incubated with PBS or WGA conjugated with Alexa488 (1:1000; Cat. W11261; Molecular Probes) or Alexa594 (1:1000; Cat. W11262; Molecular Probes) to label the plasma membrane. After washing, sections were incubated with permeabilization buffer (0.4% Triton X-100 and 2% FBS in PBS) for 1 h, and then in PBS with rabbit monoclonal anti-MOR antibody (1:500; Cat. NBP1-96656; Novus Biologicals), mouse monoclonal anti-NeuN antibody (1:1000; Cat. MAB377; Millipore), rabbit polyclonal anti-NeuN antibody (1:1000; Cat. GTX37604; GeneTex), rabbit monoclonal anti-AP2 α antibody (1:1000; Cat. GTX62588; GeneTex), or mouse monoclonal anti-clathrin heavy chain antibody (1:1000; Cat. GTX22731; GeneTex) for 24 h. Subsequently, sections were washed 4 × with the washing buffer (0.2% Triton X-100 in PBS) and then incubated in PBS containing Alexa488-conjugated goat anti-rabbit IgG antibody (1:1000; Cat. A-11008; Invitrogen), Alexa488-conjugated goat anti-mouse IgG antibody (1:1000; Cat. A-11001; Invitrogen), Alexa568-conjugated goat anti-rabbit IgG antibody (1:1000; Cat. A-11011; Invitrogen), Alexa568-conjugated goat anti-mouse IgG antibody (1:1000; Cat. A-11004; Invitrogen), or DAPI for 1 h. The slides were then washed 3 × with PBS and mounted with glycerol. Fluorescence images were captured using a laser confocal microscope (TCS SP5II, Leica) and acquired with uniform gain and exposure time. The number of puncta representing the cytoplasmic MOR (diameter > 0.3 µm) per cell was measured from confocal images of 5–10 randomly selected fields/experiment. All figures were generated using Adobe Photoshop (CS4) software. Quantification of images was performed with ImageJ (NIH) (Huang et al., 2015a).

### Internalization assay

The PathHunter GPCR internalization assay (DiscoveRx) was performed according to the manufacturer’s protocol. Briefly, we used PathHunter U2OS-MOR, U2OS-DOR, and U2OS-KOR cells with complementary pieces of β-galactosidase genetically fused to the receptor and a component of the endocytotic vesicle, respectively. In the present study, when activated opioid receptor interacts with endosomes, the two fusion proteins form a complete enzyme whose activity is detected by chemiluminescence. Cells were grown to confluence in McCoy’s 5A medium (GIBCO) containing 10% FBS, 100 units/mL penicillin, 100 μg/mL streptomycin, 20 μg/mL G418 (Sigma), 5 μg/mL hygromycin B (InvivoGen) and 25 mM HEPES in T-175 tissue-culture flasks (Corning). Cells were harvested with Cell Detachment Reagent (DiscoveRx), seeded into black, 384-well assay plates (Corning) with CP5 reagent (DiscoveRx) at 5,000 cells per well, and incubated for 24 h before experiments. For high-throughput screening, cells were treated with 5 µL of HBSS, containing either morphine alone or morphine in the presence of compounds (∼5 mM in each well) from natural-compound library plates (Greenpharma), at final concentrations of 0.3 µM and 5 µM, respectively. The potency and efficacy of compounds in promoting MOR endocytosis were determined in the absence or presence of the final concentration of morphine (0.3 µM; ∼EC10). Cells were incubated at room temperature for 1.5 h, followed by addition of 8 µL of PathHunter Detection kit (DiscoveRx) for 1 h, and analyzed for chemiluminescence on a Victor 2 plate reader (PerkinElmer). The experiments used to make each figure were run on the same day and using the same generation of cells to ensure best comparability of the data.

### β-arrestin-2 recruitment (β-arrestin assay)

The PathHunter GPCR β-arrestin-2 assay (DiscoveRx) was performed according to the manufacturer’s protocol. Briefly, when β-arrestin-2 translocates to the active receptor, complementary β-galactosidase fragments fused to the receptor and β-arrestin-2 interact to form a functional enzyme, which is detected by chemiluminescence. PathHunter CHO-K1-MOR cells were grown to confluence in F12 medium (GIBCO) containing 10% FBS, 100 units/mL penicillin, 100 µg/mL streptomycin, 200 μg/mL G418, and 20 µg/mL hygromycin B in T-175 tissue culture flasks and harvested with Cell Detachment Reagent. Cells (5,000 cells per well) were then seeded into black, 384-well assay plates with CP2 reagent (DiscoveRx) and incubated for 24 h before experiments. The effects of compounds on opioid-mediated β-arrestin-2 recruitment were determined by treating cells with 5 µL of HBSS, either containing various concentrations of morphine alone or in the presence of the compounds, and incubated at room temperature for 1.5 h. Incubations were terminated by the addition of 8 µL of PathHunter Detection kit for 1 h. Luminescence was detected using a Victor 2 plate reader. Experiments were run on the same day and using the same generation of CHO-K1-MOR cells to ensure best comparability of the data.

### cAMP accumulation (cAMP assay)

HEK-MOR cells (provided by Dr. Ping-Yee Law, University of Minnesota, USA) were cultured in DMEM (GIBCO) supplemented with 10% FBS, 100 units/mL penicillin, 100 µg/mL streptomycin, 400 µg/mL G418, and 2 mM L-glutamine in T-175 tissue culture flasks and harvested with trypsin/EDTA solution (GIBCO). Cells were plated at 72,000 per well under 100 µL/well of DMEM in 96-well, solid-bottom, white plates (GIBCO) and under 50 µL/well of HBSS containing forskolin or 3-isobutyl-1-methylxanthine at final concentrations of 1 µM and 500 µM, respectively. After 30 min of incubation at room temperature, the concentration of cAMP was determined using a LANCE Ultra cAMP Assay kit (Perkin Elmer). Two hours later, plate fluorescence was measured using a Victor 2 plate reader with excitation at 330 nm and emission at 615 nm and 665 nm. The effects of compounds on MOR-mediated inhibition of cAMP production were determined by treating cells with various concentrations of morphine alone or in the presence of the compounds. The results were presented as percent inhibition of forskolin-stimulated cAMP accumulation: [1 - (cAMP compounds/forskolin/cAMPforskolin)] × 100 %

### Membrane potential assay

Mouse pituitary AtT-20 cells (ATCC) endogenously expressing GIRK1/GIRK2 channels were cultured in T-175 tissue culture flasks in DMEM containing 10% FBS, 100 units/mL penicillin, and 100 μg/mL streptomycin and were harvested into trypsin/EDTA solution. Cells at 25,000 per well were transiently transfected with *myc*-tagged human MOR plasmid (provided by Dr. Ping-Yee Law, University of Minnesota, USA) or mutants using the NEPA21 electroporator gene transfection system (Nepa Gene) and were subsequently seeded into black, 96-well assay plates with clear, flat bottoms (Corning). The poring pulse conditions for electroporation were as follows: 110 V, pulse length 7.5 ms, inter-pulse interval 50 ms, decay rate 10%, and polarity plus. The transfer pulse conditions were as follows: 20 V, pulse length 50 ms, pulse interval 50 ms, decay rate 40%, and polarity plus and minus. After 24 h, cells were serum starved for a further 3 h. This was followed by detection of potassium conductance changes using a membrane potential assay according to the manufacturer’s instructions (Molecular Devices). Briefly, cells were treated with blue membrane-potential dye for 0.5 h at 25°C, and then the effects of vehicle, convallatoxin (1 µM), morphine (10 µM), and morphine (10 µM) + convallatoxin (1 µM) on the membrane potential were evaluated in a FlexStation 3 benchtop multi-mode microplate reader (Molecular Devices). The fluorescence signal (excitation: 485 nm, emission: 525 nm) was monitored at intervals of 1.52 s, up to 150 s after treatment.

### Molecular dynamics (MD) simulation

The MD simulation was carried out to determine the natural conformation of the MOR, using the GROMACS v5.1.2 application (Berendsen et al., 1995). The force field for the whole system was GROMOS 43a1 (Chiu et al., 2009). The protein was restrained in a box of cubic shape whose edges were placed 1 nm from the complex, and an extended simple point charge water model was formed. The system was electrically neutralized by adding 20 Cl^-^ ions. Two-step energy minimization was performed using steepest descent and conjugate gradient algorithms to converge the system up to 10 kJ mol^-^1 nm^-^1. After a short energy minimization step, the system was subjected to NVT (300 K) and NPT (1 bar) equilibration with 100 ps running, and a LINCS algorithm was used to constrain the hydrogen bond lengths (Hess et al., 1997). The time step for the simulation was 2 fs. A cut-off distance of 10 Å was used for all short-range non-bonded interactions and a 12 Å Fourier grid spacing was used in the particle mesh Ewald method for long-range electrostatics. Finally, the restraints of the complex structure were removed and a 10-ns MD calculation was performed. The equilibrium structure was used to identify the binding site region using the Discovery Studio 2016/Define protocol and the binding site editing program (BIOVIA, Inc., San Diego, CA, USA).

### Spinal cord surgery

Mice were first anesthetized with ketamine/xylazine i.p. (Sigma). To express copGFP-tagged MOR or mutants, or to reduce *clathrin* and *AP2* expression in the spinal cord and DRG neurons (Wang et al., 2005), MOR expression vectors, or a shRNA against clathrin or AP2, or the universal negative control was dissolved in artificial cerebrospinal fluid, and then injected i.t. into adult mouse spinal cord using Micro-Renathane implantation tubing (Braintree Scientific) inserted into the T11–T13 intervertebral disc. After injection, the NEPA21 electroporator gene transfection system (Nepa Gene) was used to deliver plasmid or shRNA into cells through needle electrodes inserted between the L1 and L6 vertebrae. The poring pulse conditions for electroporation were as follows: 150 V, pulse length 5 ms, inter-pulse interval 50 ms, decay rate 10%, and polarity plus. The transfer pulse conditions were as follows: 20 V, pulse length 50 ms, pulse interval 50 ms, decay rate 40%, and polarity plus and minus (Lee et al., 2014). All behavioral tests were conducted at least 7 d after surgery. We confirmed that mice that received the shRNA universal negative control had no significant tissue injury or behavioral dysfunction.

### Tail-flick test

Drug-induced antinociception was evaluated using the Tail-Flick Analgesia Meter (Columbia Instruments). The basal tail-flick latency was recorded before treatment and test latencies were recorded 45, 90, and 180 min after administration of drugs. A cutoff time of 10 s was set to avoid tissue damage. The antinociceptive effect was defined as the difference between the test latency and the basal latency (test latency - basal latency) at each time point. The AUC value was obtained by calculating the area under the time-response curve of the antinociceptive effect after treatment with the drugs. The percentage of the maximum possible effect (percentage of MPE) was calculated according to the following equation: percentage of MPE = [(test latency - baseline latency) ÷ (cutoff time - baseline latency)] × 100. The ED50 was determined by the up-and-down method (Crocker and Russell, 1984). Briefly, a series of test levels was chosen with equal spacing between each log of drug dose. Then, a series of trials (n ≧ 6) was performed following the rule that the dose is reduced on the next trial if inhibition of the tail-flick response is observed and increased if no inhibition is observed. Each mouse underwent only one trial. The ED50 value was derived from the equation ED50 = Xf + k × d, where Xf was the last dose administered, k was the tabular value, and d was the interval between doses.

To determine the chronic effects of compounds on the development of morphine tolerance, mice were chronically injected s.c. with vehicle, morphine (5 or 10 mg/kg), convallatoxin (0.5 or 1 mg/kg), or morphine (5 or 10 mg/kg) + convallatoxin (0.5 mg/kg) twice daily for 8 d. The antinociceptive effect of each treatment was determined as the radiant-heat tail-flick latency on d 1, and again after the final treatment on d 8. To determine the acute effects of compounds on morphine tolerance, mice were first injected s.c. with morphine at 10 mg/kg, twice daily for 8 d to generate morphine-tolerant mice. At 12 h after the last morphine treatment, mice were administered s.c. with acute vehicle, morphine (10 mg/kg), convallatoxin (1 mg/kg), or morphine (10 mg/kg) + convallatoxin (0.5 or 1 mg/kg) and the radiant-heat tail-flick latency was determined at the indicated time points.

### Open-field test

Testing was carried out in an open-field chamber 42 × 42 cm, 35 cm in height, under 120-lx lighting as previously described (Wu et al., 2010). Immediately after administration of morphine at 10 mg/kg, convallatoxin at 0.5 mg/kg, or vehicle, the mice were placed in the open-field apparatus along the wall, and behavior recorded with beam-break technology for 30 min. Data were analyzed with Prism v. 5.0 software (GraphPad).

### CFA-induced chronic inflammatory pain and Von Frey hair test

Chronic inflammatory pain was induced by intraplantar injection of CFA. Briefly, each B6 mouse was injected with 20 µL of either saline or CFA in the footpad of the right hind paw under isoflurane anesthesia on PID 0. From PID 14 to 18, saline- and CFA-injected mice were administered vehicle, morphine (10 mg/kg), convallatoxin (0.5 mg/kg), or morphine (10 mg/kg) + convallatoxin (0.5 mg/kg), s.c., twice daily. Mechanical allodynia was subsequently evaluated using von Frey filaments (range 0.1–1 g; IITC Life Science). Mice were placed on a mesh floor with 5 × 5 mm holes, covered with a cup to prevent visual stimulation, and allowed to adapt for 1 h prior to testing. On test days, withdrawal responses following hind paw stimulation were measured, with filaments applied to the middle of the plantar surface of the hind paw. Mechanical allodynia was defined as change in the pressure required to induce withdrawal. *Statistics.* Generally, in the present study, cell lines were not validated by genomic testing but had previously been tested for mycoplasma contamination. All mouse and in vitro experiments were repeated multiple times as indicated in the figures and figure legends. No statistical method was used to predetermine sample size. There were no exclusion criteria. No samples, mice, or data points were excluded from the reported analyses. No samples or mice were randomly assigned to experimental groups. Investigators were blinded to group allocation during the experiment and when assessing the outcome. Data are presented as the mean ± SEM. All data were tested for normal distribution and equal variance between groups using Prism v. 5.0 software (GraphPad). Multiple groups were compared using 1-way ANOVA with Newman-Keul’s post hoc test or 2-way ANOVA with Bonferroni’s post hoc test, whereas pairs of groups were compared using the two-sided Student’s *t*-test in Prism v. 5.0 software (GraphPad). *p ≤* 0.05 was considered statistically significant.

### Study approval

The protocol was approved by the Institutional Animal Care and Use Committee of the National Health Research Institutes, Taiwan. Animal experiments were conducted in accordance with the Policies on the Use of Animals in Neuroscience Research and the ethical guidelines for investigations of experimental pain in conscious animals established by the International Association for the Study of Pain.

## Author contributions

P.K.C. performed the in vivo experiments, interpreted the data, and wrote the manuscript. Y.Y.K. performed the computer modeling. H.F.C., Y.H.H., and S.C.C. performed the in vitro experiments. L.C.O. provided advice on the high-throughput screen and in vitro experiments and wrote the manuscript. W.T.C provided MOR-KO mice. Y.C.L., J.Y.C., P.T.L., S.H.U., J.T.A.H., and C.S. advised on the project. P.L.T. provided advice on the in vivo experiments. P.Y.L. provided advice on the in vitro experiments. H.H.L. provided advice on the in vivo experiments and reviewed the manuscript.S.H.Y. conceived and supervised the project, designed the experiments, performed the in vivo experiments, interpreted the data, and wrote the manuscript.

## Acknowledgments

This work was supported by grants 103-2325-B-400-018, 104-2325-B-400-008, 105-2325-B-400-007, and 101-2311-B-400-005-MY3 from the Ministry of Science and Technology, and Intramural Research Program of the National Health Research Institutes, Taiwan.

## Competing financial interests

The authors have declared that no conflict of interest exists.

## FIGURES AND FIGURE LEGENDS

**Figure 1-Supplementary Table 1.** MOR functional analysis for combinations of morphine and CTSs. The potency and efficacy of morphine in the absence and presence of each CTS are determined using the MOR endocytosis assay, the cAMP assay, and the β-arrestin assay, respectively. The efficacy of each treatment is calculated as a percentage of the Emax to morphine (*n* =5). All results are mean ± SEM.

| | MOR endocytosis assay | | β-arrestin assay | | cAMP assay | |
| --- | --- | --- | --- | --- | --- | --- |
|  | Morphine EC<sub>50</sub> | Efficacy | Morphine EC<sub>50</sub> | Efficacy | Morphine EC<sub>50</sub> | Efficacy |
| Vehicle control | 0.47 ± 0.05 μM | 100 ± 5.4 % | 0.18 ± 0.04 μM | 100 ± 2.6 % | 2.65 ± 4.87 nM | 100 ± 5.5 % |
| Gitoxigenin (1 μM) | 0.61 ± 0.26 μM | 286.3 ± 2.1 % | 0.17 ± 0.04 μM | 99.2 ± 5.2 % | 4.95 ± 1.81 nM | 105.8 ± 1.4 % |
| Bufalin (1 μM) | 0.6 ± 0.38 μM | 376 ± 1.3 % | 0.22 ± 0.05 μM | 103.5 ± 4.3 % | 11.02 ± 1.75 nM | 105.9 ± 0.6 % |
| Convallatoxin (1 μM) | 0.46 ± 0.26 μM | 356.4 ± 8.6 % | 0.20 ± 0.14 μM | 99.2 ± 3.7 % | 8.09 ± 0.1 nM | 105.1 ± 0.9 % |

**Figure 1-Supplementary Figure 1.**
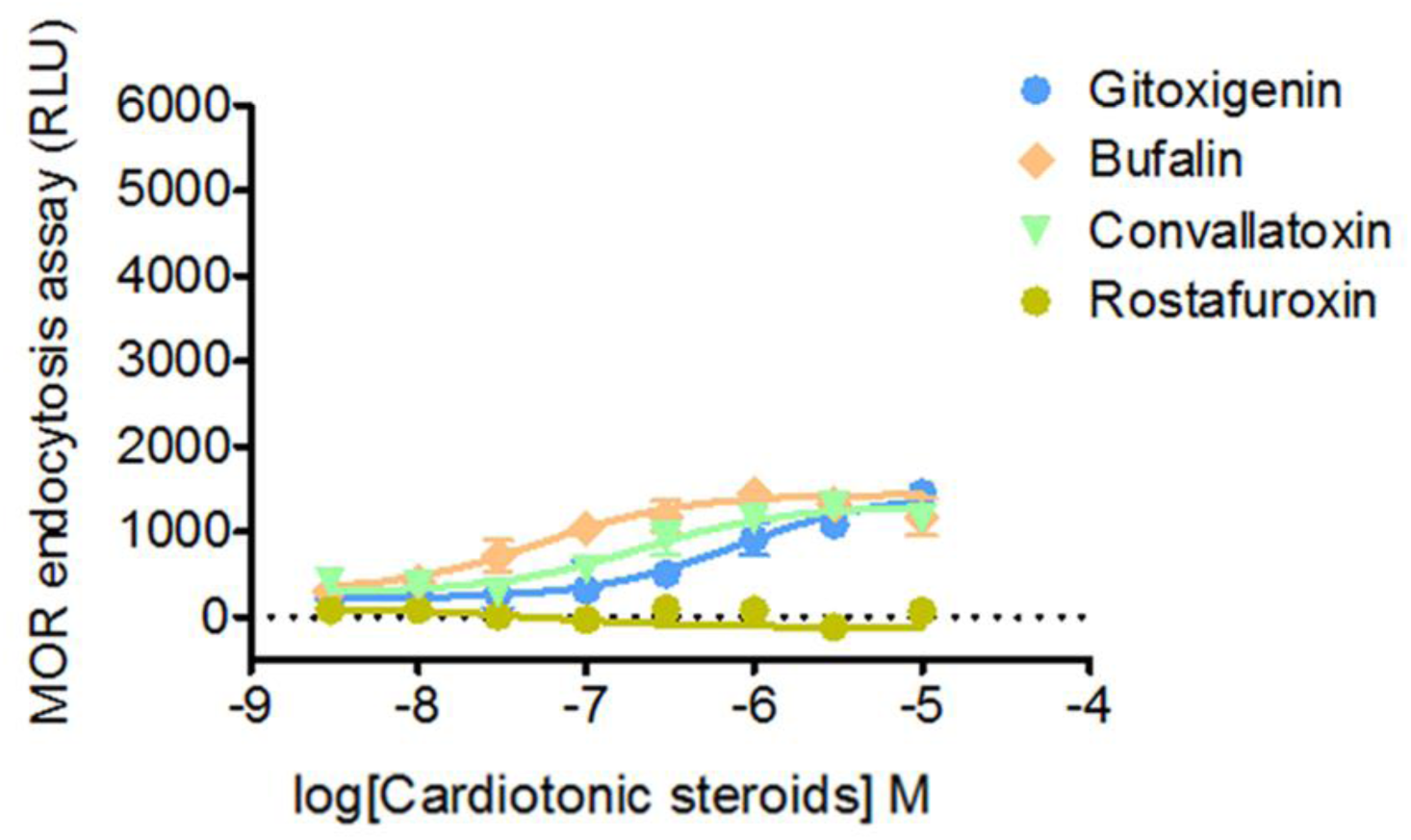
Effect of cardiotonic steroids (CTSs) on mu-opioid receptor (MOR) endocytosis. MOR endocytosis is only slightly upregulated by CTSs in the absence of morphine. The effects of CTSs on MOR endocytosis are determined by treating U2OS-MOR cells with various concentrations of gitoxigenin, bufalin, convallatoxin, and rostafuroxin (*n* = 5). Values are means ± SEM (all figures).

**Figure 1-Supplementary Figure 2.**
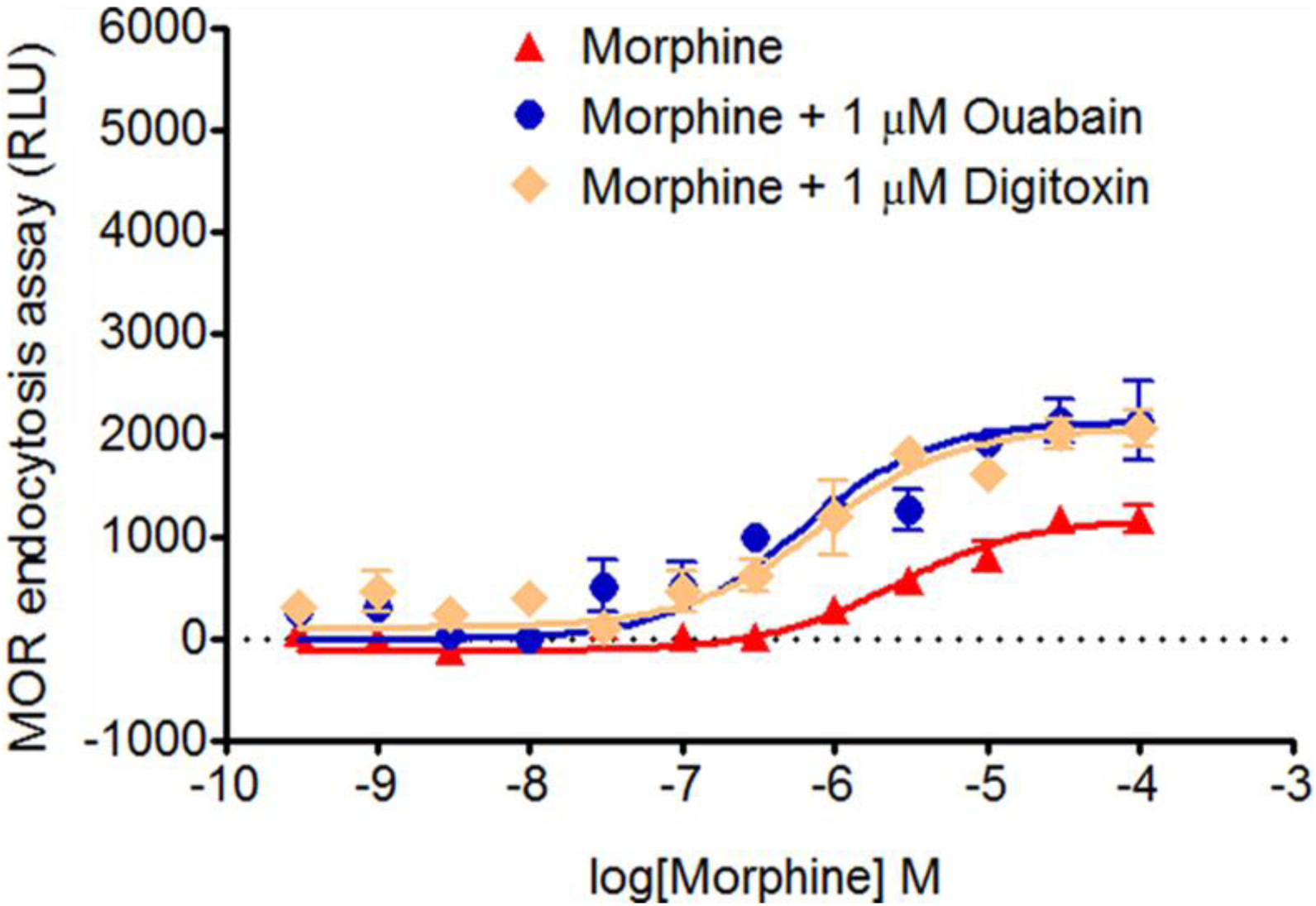
Effect of ouabain and digitoxin on morphine-induced MOR endocytosis. Morphine-induced MOR endocytosis is enhanced by CTSs. The effects of CTSs on opioid-induced MOR endocytosis are determined by treating U2OS-MOR cells with various concentrations of morphine alone, or with morphine in the presence of ouabain or digitoxin (1 µM; *n* = 5).

**Figure 1-Supplementary Figure 3.**
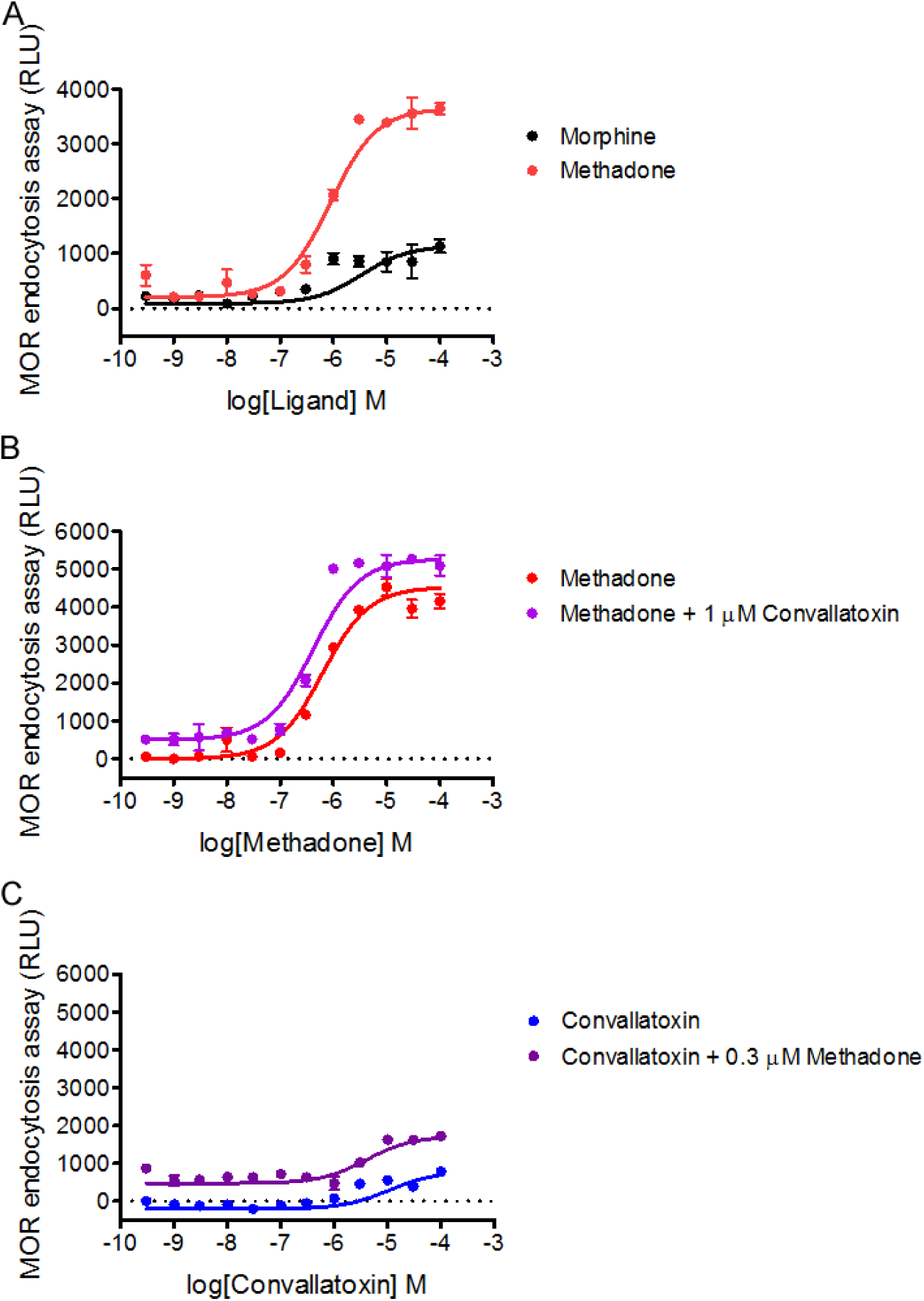
Effect of convallatoxin on methadone-induced MOR endocytosis. (**A**) Concentration-response curves of MOR endocytosis against morphine or methadone for (*n* = 5). (**B**) Methadone induction of MOR endocytosis in the absence or presence of convallatoxin (1 µM; *n* = 5). (**C**) Convallatoxin induction of MOR endocytosis in the absence or presence of methadone (0.3 µM; *n* = 5). For data in (**A**–**C**), MOR internalization is measured by an enzyme complementation assay in U2OS-MOR cells.

**Figure 2-Supplementary Figure 1.**
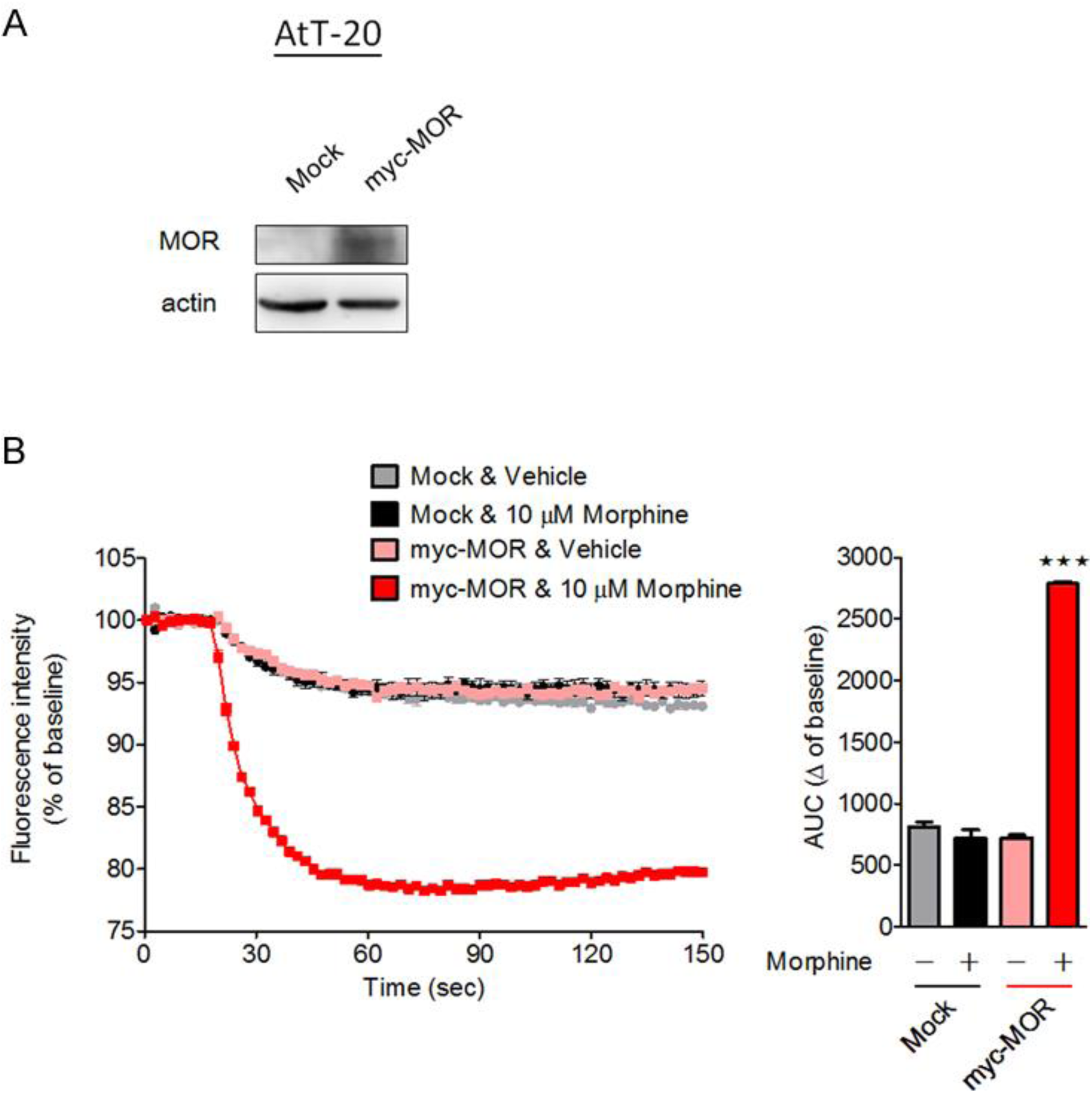
Morphine-induced membrane hyperpolarization is dependent on MOR expression in AtT-20 cells. (**A**) Mouse pituitary AtT-20 cells are transiently transfected with vehicle or the *myc-MOR* expression plasmid 24 h prior to the experiment. The expression of *myc-MOR* in mock- or *myc-MOR*-transfected cells is confirmed by immunoblotting. (**B**) The effects of morphine on membrane potential are determined by comparing potentials in the presence or absence of morphine at 10 µM. The calculated area under the curve (AUC) represents the total drug exposure integrated over time. *F*_3,16_ = 614; *p* < 0.001 (1-way ANOVA). *n* = 5 per group. ***, *p <* 0.001 versus mock + vehicle control group (Newman-Keul’s post hoc test).

**Figure 2-Supplementary Figure 2.**
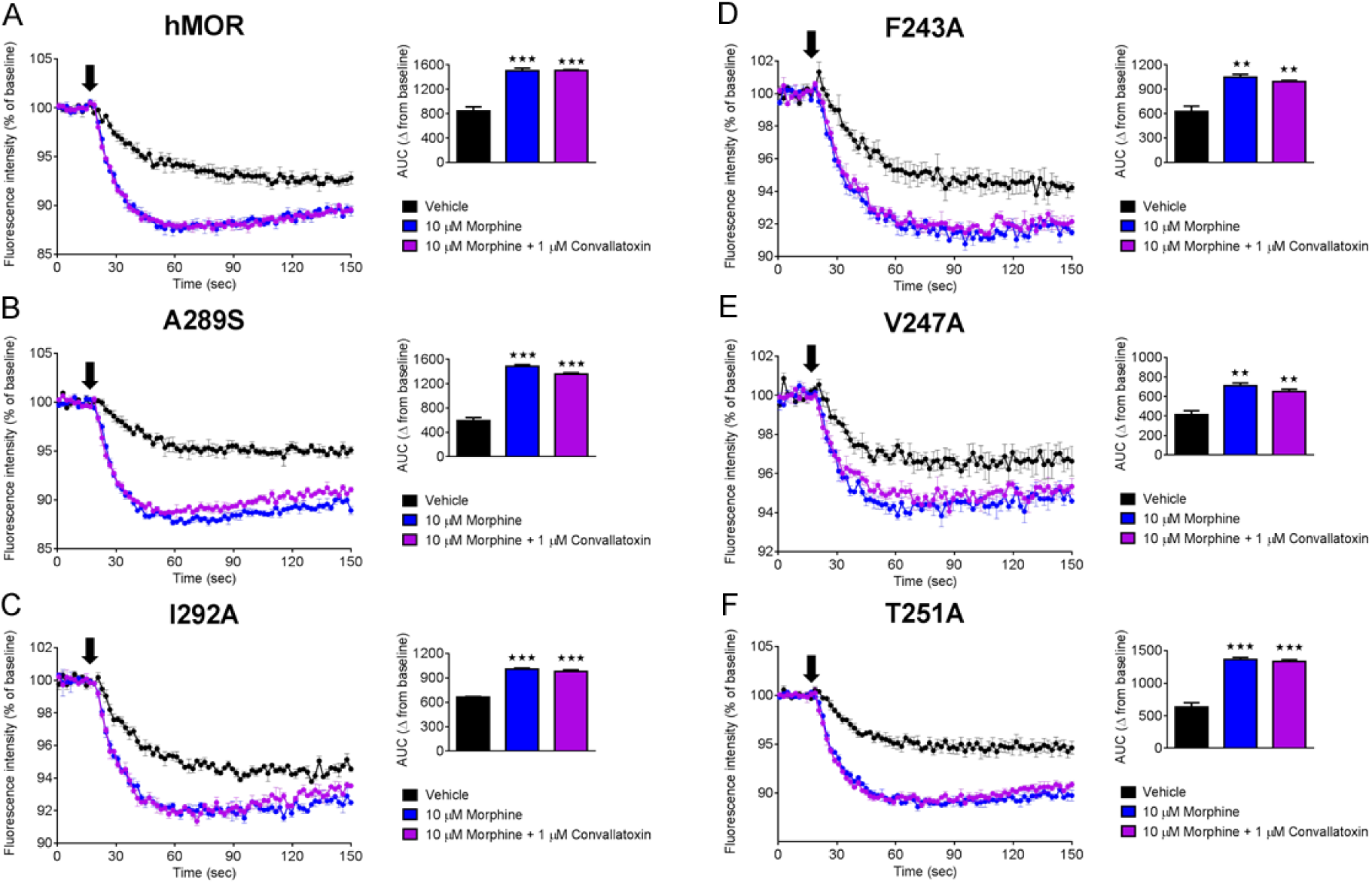
Membrane potential analysis of mutants in the presence of morphine and morphine + convallatoxin. AtT-20 cells are transfected with plasmid expressing *myc-MOR* or mutants 24 h prior to any membrane potential assay. (**A**–**F**) Effects of morphine and morphine + convallatoxin on AtT-20 cells expressing (**A**) human MOR, or mutated variants of it, namely (**B**) A289S, (**C**) I292A, (**D**) F243A, (**E**) V247A, and (**F**) T251A, all measured using a membrane potential assay. (**A**) MOR AUC: *F*_2,6_ = 69.09; *p <* 0.0001, (**B**) A289S AUC: *F*_2,6_ = 179.5; *p <* 0.0001, (**C**) I292A AUC: *F*_2,6_ = 167.9; *p <* 0.0001, (**D**) F243A AUC: *F*_2,6_ = 28.43; *p <* 0.001, (**E**) V247A AUC: *F*_2,6_ = 25.62; *p <* 0.01, and (**F**) T251A AUC: *F*_2,6_ = 85.48; *p <* 0.0001 (1-way ANOVA). **, *p* <0.01; ***, *p* < 0.001 versus vehicle control group (Newman-Keul’s post hoc test). All *n* = 3 per group.

**Figure 3-Supplementary Figure 1.**
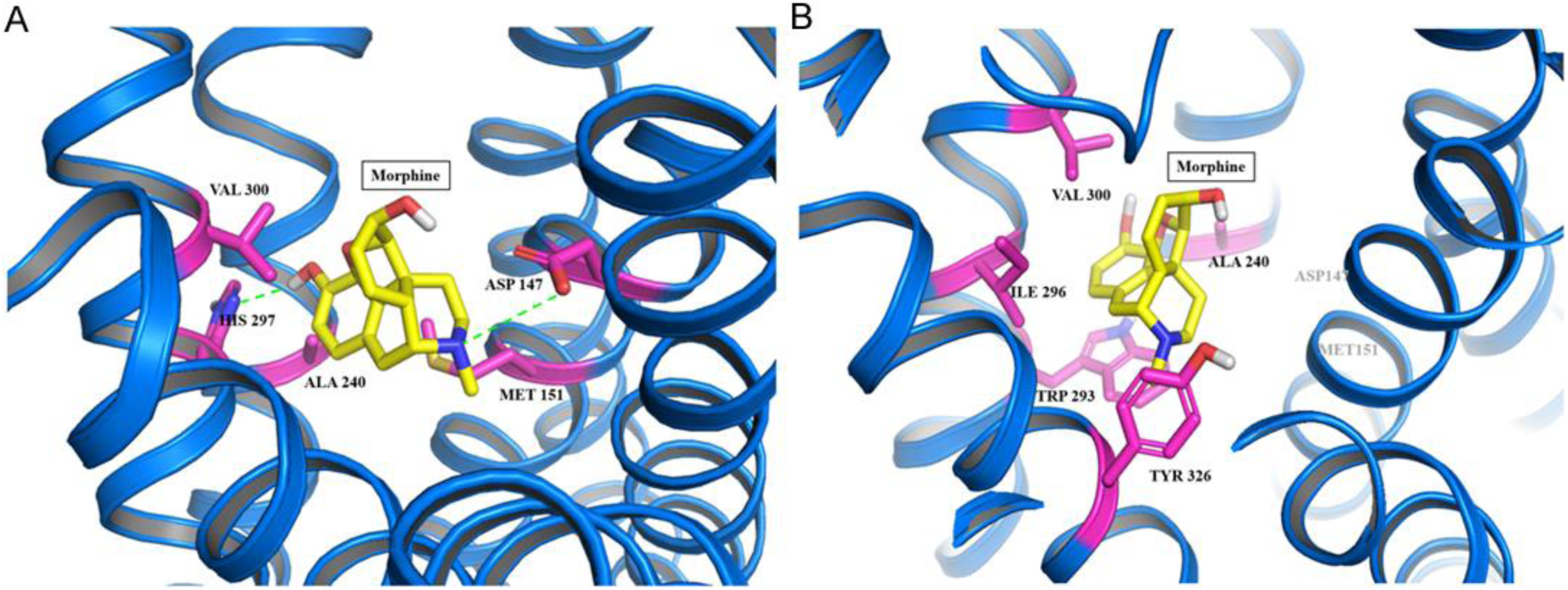
Molecular-dynamics analysis of the MOR-morphine complex. (**A**) The binding analysis for morphine docking to mouse MOR. (**B**) The binding analysis for the morphine-MOR complex after 20 ns of equilibrium molecular dynamics.

**Figure 3-Supplementary Table 1.** Top residues of mouse MOR with higher interactions with convallatoxin. According to the Discovery Studio 2016/ Analyze Ligand Poses program analysis, different docking-ligand poses will generate different interactions with the binding site residues. The residues with the most energetically favorable interactions between convallatoxin and mouse MOR are illustrated in this table. Thr249 and Ile290 support a hydrogen bond with convallatoxin. The F241, I290, V245, and A287 residues contribute hydrophobic interactions with the compound.

| Top Residues with Favorable Interactions |  |  |  |  |  |  |  |
| --- | --- | --- | --- | --- | --- | --- | --- |
| Residue | Favorable | Unfavorable | HydrogenBond | Charge | Hydrophobic | Halogen | Other |
| A:F241 | 40 | 0 | 0 | 0 | 40 | 0 | 0 |
| A:I290 | 38 | 0 | 4 | 0 | 36 | 0 | 0 |
| A:V245 | 21 | 0 | 0 | 0 | 21 | 0 | 0 |
| A:T249 | 11 | 0 | 11 | 0 | 0 | 0 | 0 |
| S:A287 | 11 | 0 | 0 | 0 | 11 | 0 | 0 |

**Figure 3-Supplementary Figure 2.**
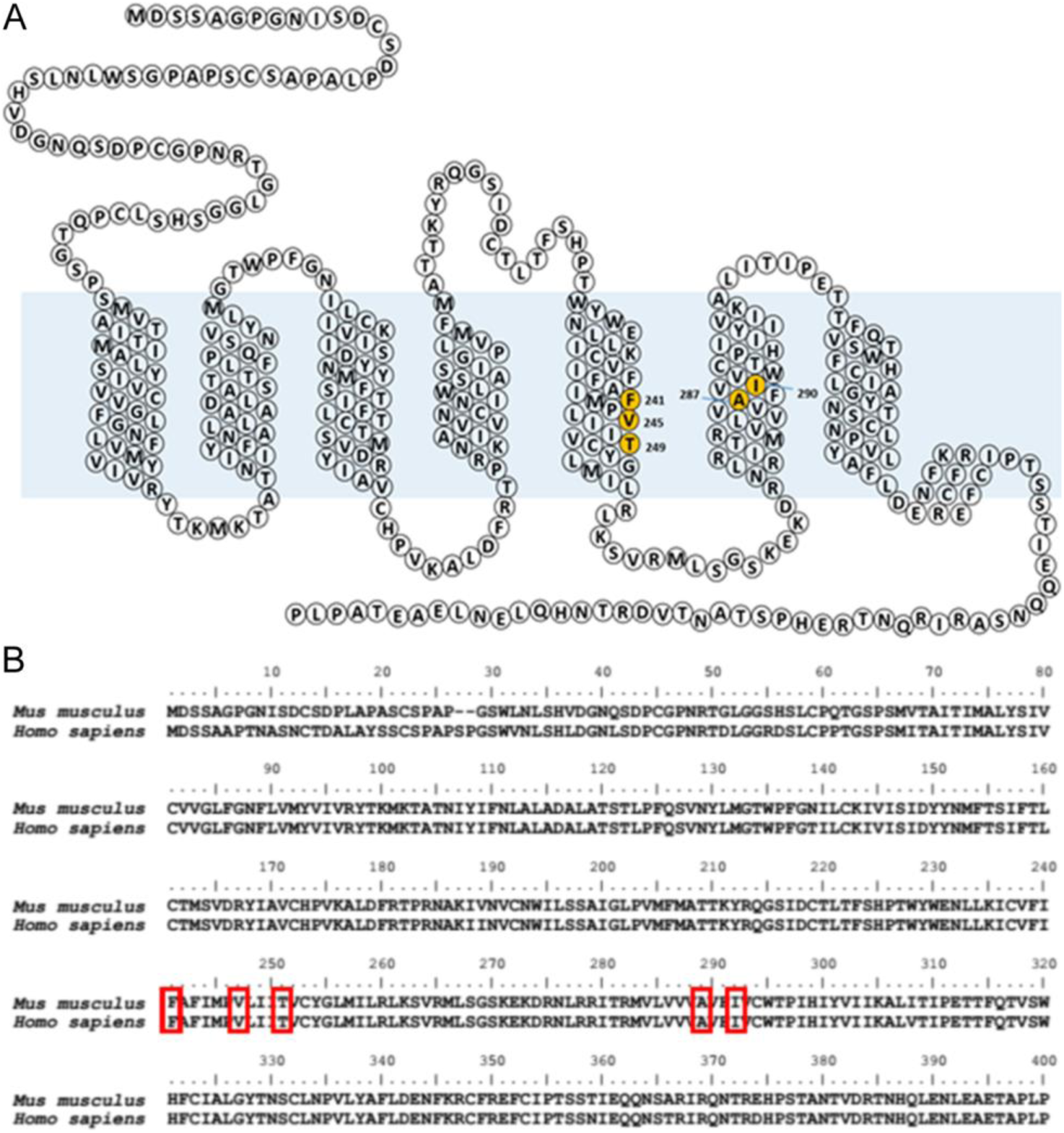
Sequencing alignment for MOR and docking analysis for convallatoxin. (**A**) Mouse MOR snake plot showing extracellular loops and transmembrane domains (upper panel). Important residues for convallatoxin binding are highlighted. (**B**) Sequence alignment of MOR from human and mouse. Red boxes indicate residues important for convallatoxin function. Human F243, V247, T251, A289, and I292 were mutated in this study.

**Figure 3-Supplementary Table 2.**
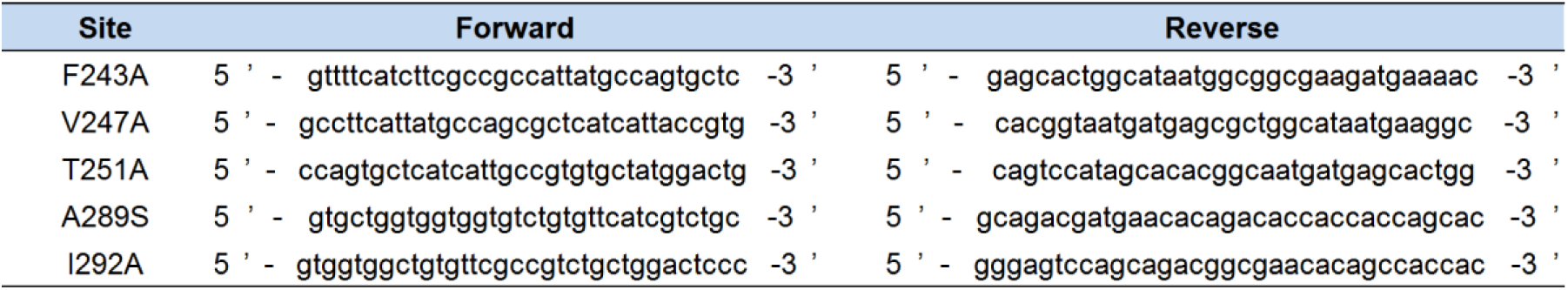
Primers used in site-directed mutagenesis.

**Figure 4-Supplementary Figure 1.**
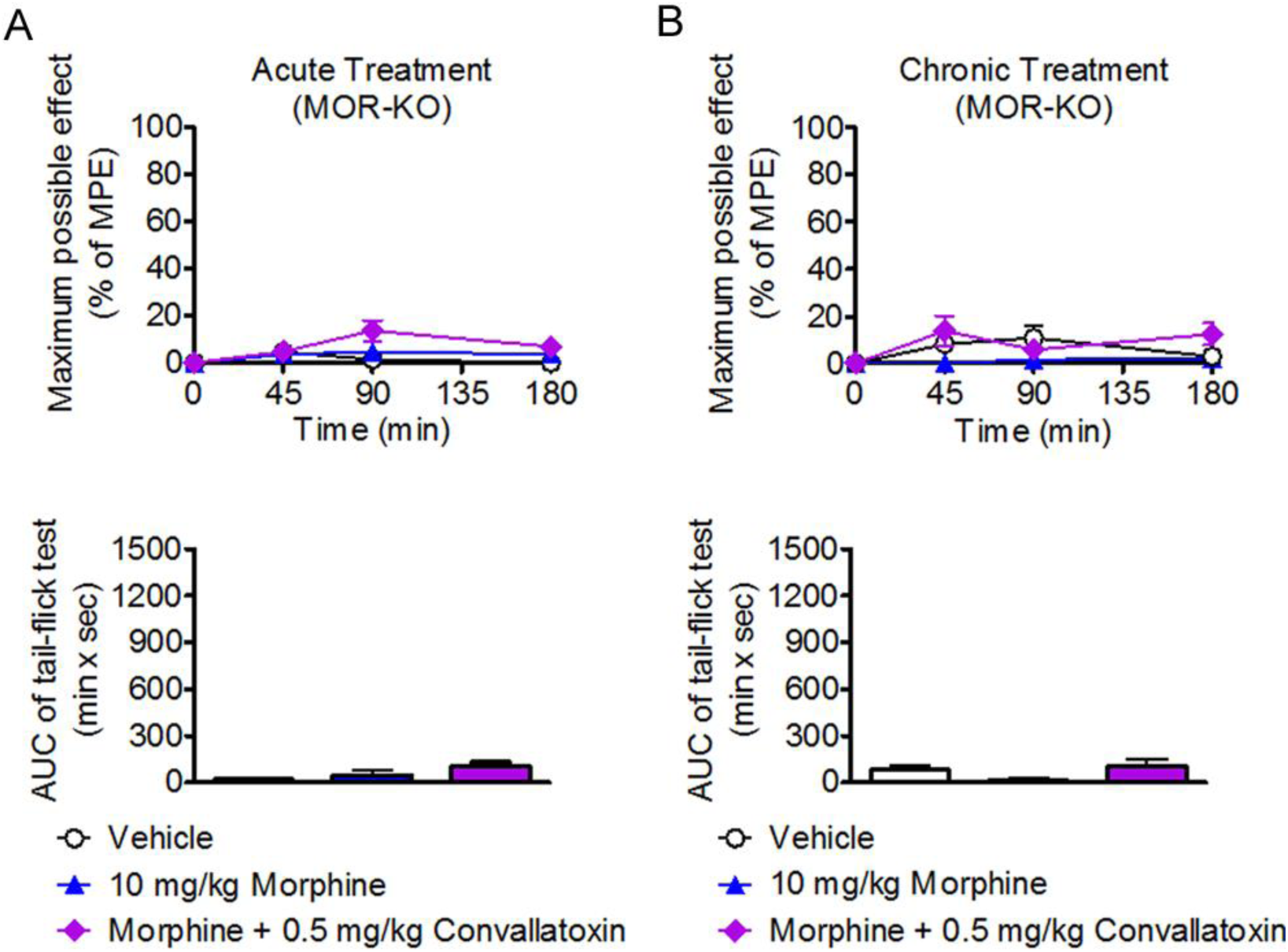
Effect of convallatoxin in morphine antinociceptive tolerance and dependence in MOR-KO mice. (**A** and **B**) MOR-KO mice are chronically injected s.c. with vehicle, morphine, or morphine + convallatoxin twice daily for 8 d. The acute antinociceptive effect of each treatment is determined by testing the tail-flick latency on d 1 (**A**), and again after the final treatment on d 8 (**B**). (**A**, upper panel) Treatment *F*_2,12_ = 1.92, time *F*_3,36_ = 5.45, interaction *F*_6,36_ = 2.47; (**B**, upper panel) Treatment *F*_2,12_ = 2.95, time *F*_3,36_ = 5.21, interaction *F*_6,36_ = 2.62; all *p* > 0.05 (2-way ANOVA). Quantitative results from (**A**) and (**B**) are presented as the AUC. (**A**, lower panel) *F*_2,12_ = 2.26; (**B**, lower panel) *F*_2,12_ = 2.54; all *p* > 0.05 (1-way ANOVA). Data in **A** and **B**, *n* = 5 per group. MPE, maximum possible effect.

**Figure 4-Supplementary Figure 2.**
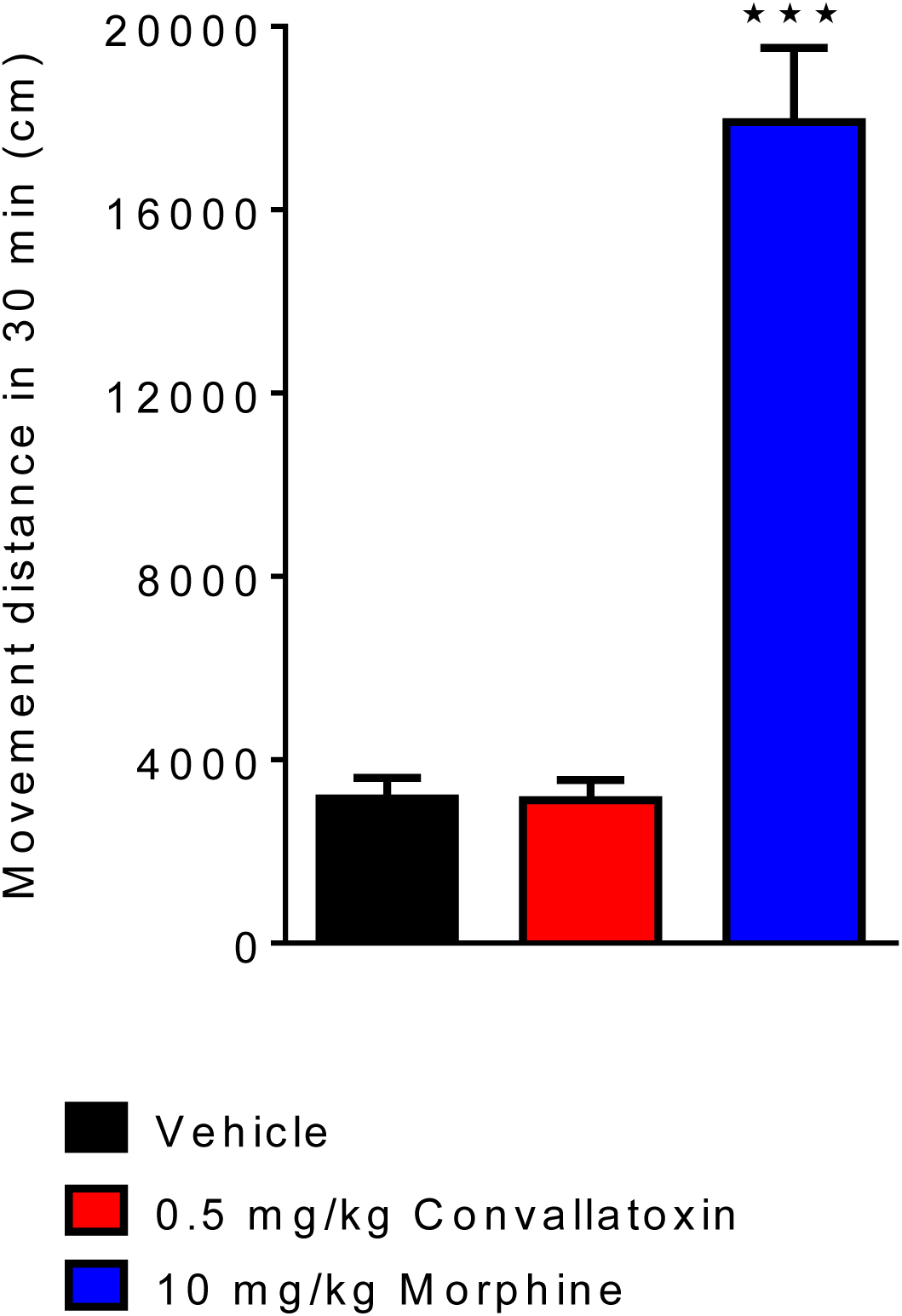
Effect of morphine or convallatoxin on locomotor activity in mice. Open-field tests are carried out in an open-field chamber (42 × 42 cm, 35 cm height) under 120-lx lighting. Morphine (10 mg/kg, s.c.) elicited hyper-locomotion activity as indicated by an increased total distance walked. Treatment with convallatoxin alone did not change the total distance compared with vehicle-treated group. *F*_2,12_ = 72.67; *p* < 0.0001 (1-way ANOVA). *n* = 5 per group. ***, *p <* 0.001 versus vehicle-treated group (Newman-Keul’s post hoc test).

**Figure 5-Supplementary Figure 1.**
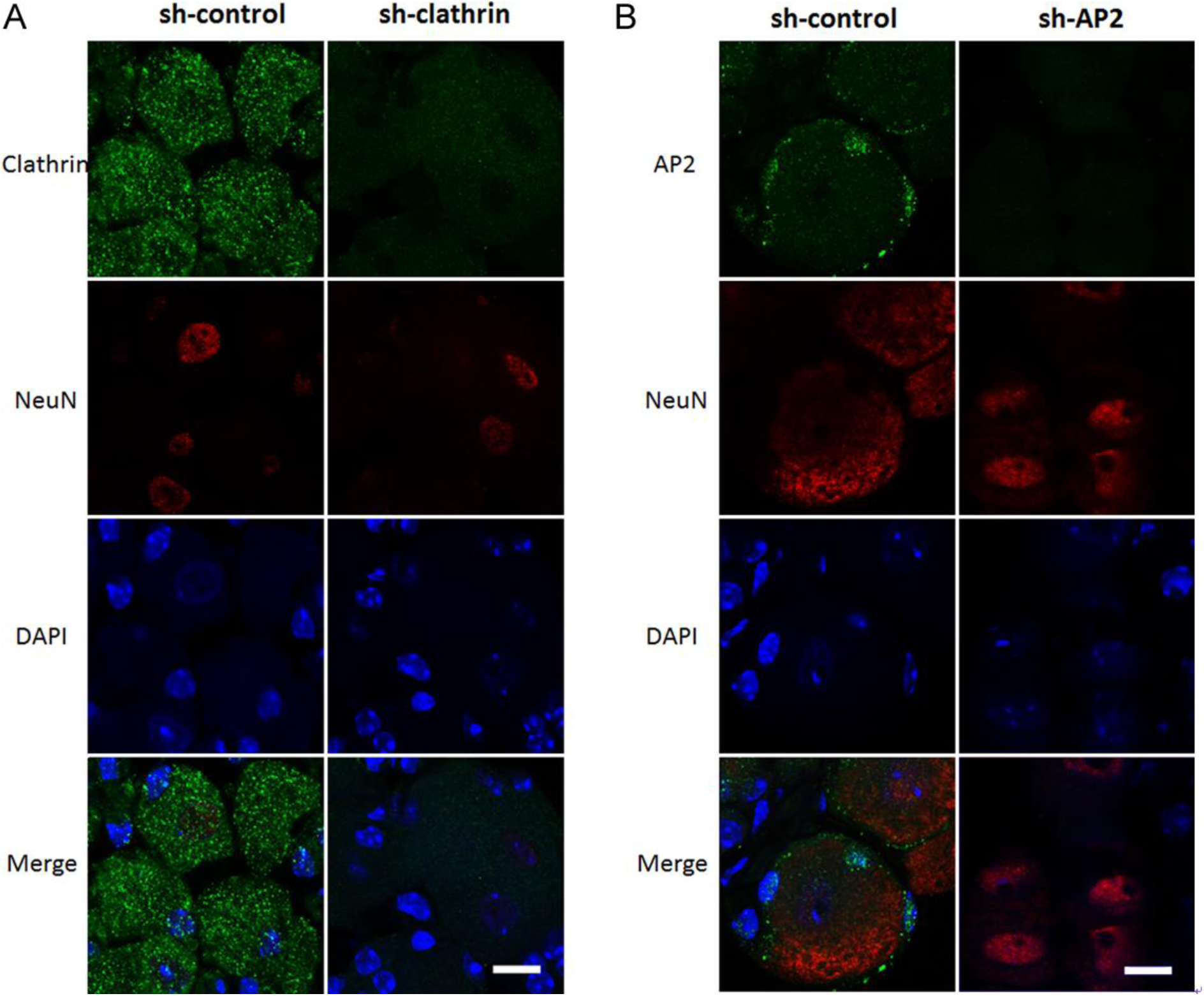
Down-regulation of *clathrin* or *AP2* in mouse DRG neurons. (**A** and **B**) shRNA (*sh-control*, *sh-clathrin,* or *sh-AP2*) is transfected into the spinal cord of B6 mice by in vivo electroporation. Seven days later, mice are sacrificed and the expression levels of *clathrin* (**A**, green), *AP2* (**B**, green) and *NeuN* (**A** and **B**, red) are visualized by immunostaining. DAPI (blue) is a nuclear marker. *n* = 5–10 per group. Scale bars, 20 µm.

**Figure 6-Supplementary Figure 1.**
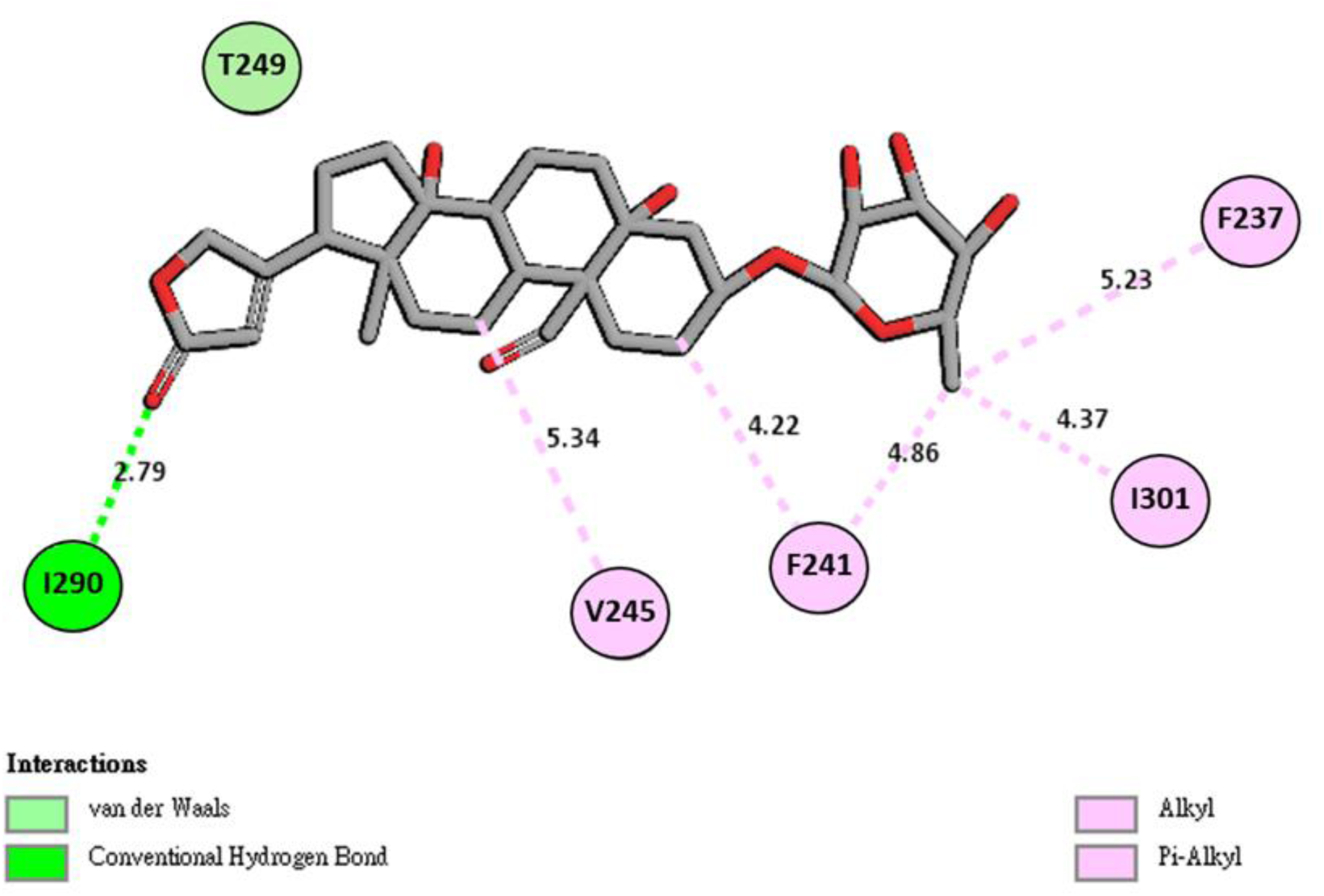
Two-dimensional representation of the ligand-mouse MOR interaction. The 2D analysis was run with the following default settings: Hydrogen bond inferred below 3.4 Å, alkyl interaction inferred below 5.5 Å. After the docking and MD simulations, I290 forms a hydrogen bond with the carbonyl group of the furanone moiety of convallatoxin, and F241 forms pi-alkyl hydrophobic interactions with the methyl substituent on the tetrahydropyrantriol moiety, and with the methylene group of the second carbon of convallatoxin. I301 can form an alkyl-alkyl hydrophobic interaction with the methyl substituent of the tetrahydropyrantriol moiety.

## References

Al-Hasani, R., and M.R. Bruchas. 2011. Molecular Mechanisms of Opioid Receptor-Dependent Signaling and Behavior. Anesthesiology 115:1363–1381.

Alvarez, V.A., S. Arttamangkul, V. Dang, A. Salem, J.L. Whistler, M. von Zastrow, D.K. Grandy, and J.T. Williams. 2002. µ-Opioid Receptors: Ligand-Dependent Activation of Potassium Conductance, Desensitization, and Internalization. The Journal of Neuroscience 22:5769–5776.

Becker, D.E. 2010. Pain Management: Part 1: Managing Acute and Postoperative Dental Pain. Anesthesia Progress 57:67–79.

Berendsen, H.J.C., D. van der Spoel, and R. van Drunen. 1995. GROMACS: A message-passing parallel molecular dynamics implementation. Computer Physics Communications 91:43–56.

Blanchet, C., M. Sollini, and C. Lüscher. 2003. Two distinct forms of desensitization of G-protein coupled inwardly rectifying potassium currents evoked by alkaloid and peptide µ-opioid receptor agonists. Molecular and Cellular Neuroscience 24:517–523.

Burford, N.T., M.J. Clark, T.S. Wehrman, S.W. Gerritz, M. Banks, J. O’Connell, J.R. Traynor, and A. Alt. 2013. Discovery of positive allosteric modulators and silent allosteric modulators of the µopioid receptor. Proceedings of the National Academy of Sciences of the United States of America 110:10830–10835.

Chen, S.-L., H.-I. Ma, J.-M. Han, R.-B. Lu, P.-L. Tao, P.-Y. Law, and H.H. Loh. 2010. Antinociceptive effects of morphine and naloxone in mu-opioid receptor knockout mice transfected with the MORS196A gene. Journal of Biomedical Science 17:28–28.

Chiu, S.-W., S.A. Pandit, H.L. Scott, and E. Jakobsson. 2009. An Improved United Atom Force Field for Simulation of Mixed Lipid Bilayers. The Journal of Physical Chemistry B 113:2748–2763.

Cooke, A.E., S. Oldfield, C. Krasel, S.J. Mundell, G. Henderson, and E. Kelly. 2015. Morphine-induced internalization of the L83I mutant of the rat µ-opioid receptor. British Journal of Pharmacology 172:593–605.

Crocker, A.D., and R.W. Russell. 1984. The up-and-down method for the determination of nociceptive thresholds in rats. Pharmacology Biochemistry and Behavior 21:133–136.

Davis, C.N., S.R. Bradley, H.H. Schiffer, M. Friberg, K. Koch, B.-R. Tolf, D.W. Bonhaus, and J. Lameh. 2009. Differential regulation of muscarinic M1 receptors by orthosteric and allosteric ligands. BMC Pharmacology 9:14–14.

Dhami, G.K., L.B. Dale, P.H. Anborgh, K.E. O’Connor-Halligan, R. Sterne-Marr, and S.S.G. Ferguson. 2004. G Protein-coupled Receptor Kinase 2 Regulator of G Protein Signaling Homology Domain Binds to Both Metabotropic Glutamate Receptor 1a and Gαq to Attenuate Signaling. Journal of Biological Chemistry 279:16614–16620.

Duttaroy, B.S.P.M.S.A., and P. Byron C. Yoburn. 1995. The Effect of Intrinsic Efficacy on Opioid Tolerance The Journal of the American Society of Anesthesiologists 82:1226–1236.

Enquist, J., M. Ferwerda, L. Milan-Lobo, and J.L. Whistler. 2012. Chronic Methadone Treatment Shows a Better Cost/Benefit Ratio than Chronic Morphine in Mice. The Journal of Pharmacology and Experimental Therapeutics 340:386–392.

Fernández-Dueñas, V., O. Pol, P. García-Nogales, L. Hernández, E. Planas, and M.M. Puig. 2007. Tolerance to the Antinociceptive and Antiexudative Effects of Morphine in a Murine Model of Peripheral Inflammation. Journal of Pharmacology and Experimental Therapeutics 322:360–368.

Ferrari, P., M. Ferrandi, G. Valentini, and G. Bianchi. 2006. Rostafuroxin: an ouabain antagonist that corrects renal and vascular Na+-K+- ATPase alterations in ouabain and adducin-dependent hypertension. American Journal of Physiology - Regulatory, Integrative and Comparative Physiology 290:R529–R535.

Finn, A.K., and J.L. Whistler. 2001. Endocytosis of the Mu Opioid Receptor Reduces Tolerance and a Cellular Hallmark of Opiate Withdrawal. Neuron 32:829–839.

Gonzalez, L.G., W. Masocha, C. Sánchez-Fernández, A. Agil, M. Ocaña, E. Del Pozo, and J.M. Baeyens. 2012. Changes in morphine-induced activation of cerebral Na+,K+-ATPase during morphine tolerance: Biochemical and behavioral consequences. Biochemical Pharmacology 83:1572–1581.

Goodman, O.B., J.G. Krupnick, F. Santini, V.V. Gurevich, R.B. Penn, A.W. Gagnon, J.H. Keen, and J.L. Benovic. 1996. β-Arrestin acts as a clathrin adaptor in endocytosis of the β2-adrenergic receptor. Nature 383:447–450.

Gozalpour, E., R. Greupink, A. Bilos, V. Verweij, J.J.M.W. van den Heuvel, R. Masereeuw, F.G.M. Russel, and J.B. Koenderink. 2014. Convallatoxin: A new P-glycoprotein substrate. European Journal of Pharmacology 744:18–27.

Gray, J.A., A. Bhatnagar, V.V. Gurevich, and B.L. Roth. 2003. The Interaction of a Constitutively Active Arrestin with the Arrestin-Insensitive 5-HT2A Receptor Induces Agonist-Independent Internalization. Molecular Pharmacology 63:961–972.

Grecksch, G., S. Just, C. Pierstorff, A.-K. Imhof, L. Glück, C. Doll, A. Lupp, A. Becker, T. Koch, R. Stumm, V. Höllt, and S. Schulz. 2011. Analgesic Tolerance to High-Efficacy Agonists But Not to Morphine Is Diminished in Phosphorylation-Deficient S375A µ-Opioid Receptor Knock-In Mice. The Journal of Neuroscience 31:13890–13896.

He, L., J. Fong, M. von Zastrow, and J.L. Whistler. 2002. Regulation of Opioid Receptor Trafficking and Morphine Tolerance by Receptor Oligomerization. Cell 108:271–282.

He, L., J. Kim, C. Ou, W. McFadden, R.M. van Rijn, and J.L. Whistler. 2009. Methadone Antinociception Is Dependent on Peripheral Opioid Receptors. The Journal of Pain 10:369–379.

He, L., and J.L. Whistler. 2005. An Opiate Cocktail that Reduces Morphine Tolerance and Dependence. Current Biology 15:1028–1033.

Hess, B., H. Bekker, H.J.C. Berendsen, and J.G.E.M. Fraaije. 1997. LINCS: A linear constraint solver for molecular simulations. Journal of Computational Chemistry 18:1463–1472.

Huang, S.-P., J.-Y. Chien, and R.-K. Tsai. 2015a. Ethambutol induces impaired autophagic flux and apoptosis in the rat retina. Disease Models & Mechanisms 8:977.

Huang, W., A. Manglik, A.J. Venkatakrishnan, T. Laeremans, E.N. Feinberg, A.L. Sanborn, H.E. Kato, K.E. Livingston, T.S. Thorsen, R.C. Kling, S. Granier, P. Gmeiner, S.M. Husbands, J.R. Traynor, W.I. Weis, J. Steyaert, R.O. Dror, and B.K. Kobilka. 2015b. Structural insights into µ-opioid receptor activation. Nature 524:315–321.

Huh, J.R., M.W.L. Leung, P. Huang, D.A. Ryan, M.R. Krout, R.R.V. Malapaka, J. Chow, N. Manel, M. Ciofani, S.V. Kim, A. Cuesta, F.R. Santori, J.J. Lafaille, H.E. Xu, D.Y. Gin, F. Rastinejad, and D.R. Littman. 2011. Digoxin and its derivatives suppress Th17 cell differentiation by antagonizing RORt activity. Nature 472:486–490.

Keith, D.E., S.R. Murray, P.A. Zaki, P.C. Chu, D.V. Lissin, L. Kang, C.J. Evans, and M. von Zastrow. 1996. Morphine Activates Opioid Receptors without Causing Their Rapid Internalization. Journal of Biological Chemistry 271:19021–19024.

Kim, J.A., S. Bartlett, L. He, C.K. Nielsen, A.M. Chang, V. Kharazia, M. Waldhoer, C.J. Ou, S. Taylor, M. Ferwerda, D. Cado, and J.L. Whistler. 2008. Morphine-induced endocytosis of the mu opioid receptor in a novel knock in mouse enhances analgesia and reduces tolerance and dependence. Current biology : CB 18:129–135.

Labianca, R., P. Sarzi-Puttini, S.M. Zuccaro, P. Cherubino, R. Vellucci, and D. Fornasari. 2012. Adverse effects associated with non-opioid and opioid treatment in patients with chronic pain. Clinical drug investigation 32 Suppl 1:53–63.

Lamers, W.H., P.G. Mooren, A. De Graaf, and R. Charles. 1985. Perinatal development of the liver in rat and spiny mouse. European Journal of Biochemistry 146:475–480.

Laporte, S.A., R.H. Oakley, J. Zhang, J.A. Holt, S.S.G. Ferguson, M.G. Caron, and L.S. Barak. 1999. The β2-adrenergic receptor/βarrestin complex recruits the clathrin adaptor AP-2 during endocytosis. Proceedings of the National Academy of Sciences of the United States of America 96:3712–3717.

Lee, P.-T., P.-K. Chao, L.-C. Ou, J.-Y. Chuang, Y.-C. Lin, S.-C. Chen, H.-F. Chang, P.-Y. Law, H.H. Loh, Y.-S. Chao, T.-P. Su, and S.-H. Yeh. 2014. Morphine drives internal ribosome entry site-mediated hnRNP K translation in neurons through opioid receptor-dependent signaling. Nucleic Acids Research 42:13012–13025.

Loh, H.H., H.-C. Liu, A. Cavalli, W. Yang, Y.-F. Chen, and L.-N. Wei. 1998. µ Opioid receptor knockout in mice: effects on ligand-induced analgesia and morphine lethality. Molecular Brain Research 54:321–326.

Mankovsky, T., M.E. Lynch, A.J. Clark, J. Sawynok, and M.J.L. Sullivan. 2012. Pain catastrophizing predicts poor response to topical analgesics in patients with neuropathic pain. Pain Research & Management : The Journal of the Canadian Pain Society 17:10–14.

Marker, C.L., M. Stoffel, and K. Wickman. 2004. Spinal G-Protein-Gated K+ Channels Formed by GIRK1 and GIRK2 Subunits Modulate Thermal Nociception and Contribute to Morphine Analgesia. The Journal of Neuroscience 24:2806–2812.

Martini, L., and J.L. Whistler. 2007. The role of mu opioid receptor desensitization and endocytosis in morphine tolerance and dependence. Current Opinion in Neurobiology 17:556–564.

Masocha, W., G. Horvath, A. Agil, M. Ocaña, E. Del Pozo, M. Szikszay, and J.M. Baeyens. 2003. Role of Na+,K+-ATPase in Morphine-Induced Antinociception. Journal of Pharmacology and Experimental Therapeutics 306:1122–1128.

Milan-Lobo, L., and J.L. Whistler. 2011. Heteromerization of the µ- and δ-Opioid Receptors Produces Ligand-Biased Antagonism and Alters µ-Receptor Trafficking. The Journal of Pharmacology and Experimental Therapeutics 337:868–875.

Nagakura, Y., M. Okada, A. Kohara, T. Kiso, T. Toya, A. Iwai, F. Wanibuchi, and T. Yamaguchi. 2003. Allodynia and Hyperalgesia in Adjuvant-Induced Arthritic Rats: Time Course of Progression and Efficacy of Analgesics. Journal of Pharmacology and Experimental Therapeutics 306:490–497.

Nockemann, D., M. Rouault, D. Labuz, P. Hublitz, K. McKnelly, F.C. Reis, C. Stein, and P.A. Heppenstall. 2013. The K(+) channel GIRK2 is both necessary and sufficient for peripheral opioid-mediated analgesia. EMBO Molecular Medicine 5:1263–1277.

Prassas, I., and E.P. Diamandis. 2008. Novel therapeutic applications of cardiac glycosides. Nat Rev Drug Discov 7:926–935.

Ravindranathan, A., G. Joslyn, M. Robertson, M.A. Schuckit, J.L. Whistler, and R.L. White. 2009. Functional characterization of human variants of the mu-opioid receptor gene. Proceedings of the National Academy of Sciences of the United States of America 106:10811–10816.

Ritter, S.L., and R.A. Hall. 2009. Fine-tuning of GPCR activity by receptor-interacting proteins. Nature reviews. Molecular cell biology 10:819–830.

Rodal, S.K., G. Skretting, Ø. Garred, F. Vilhardt, B. van Deurs, and K. Sandvig. 1999. Extraction of Cholesterol with Methyl-β-Cyclodextrin Perturbs Formation of Clathrin-coated Endocytic Vesicles. Molecular Biology of the Cell 10:961–974.

Rowbotham, M.C., L. Twilling, P.S. Davies, L. Reisner, K. Taylor, and D. Mohr. 2003. Oral opioid therapy for chronic peripheral and central neuropathic pain. The New England journal of medicine 348:1223–1232.

Soohoo, A.L., and M.A. Puthenveedu. 2013. Divergent modes for cargo-mediated control of clathrin-coated pit dynamics. Molecular Biology of the Cell 24:1725–1734.

Sounier, R., C. Mas, J. Steyaert, T. Laeremans, A. Manglik, W. Huang, B.K. Kobilka, H. Demene, and S. Granier. 2015. Propagation of conformational changes during [mgr]-opioid receptor activation. Nature 524:375–378.

Tao, P.L., P.Y. Law, and H.H. Loh. 2010. Search for the “ideal analgesic” in pain treatment by engineering the mu-opioid receptor. IUBMB Life 62:103–111.

van Koppen, C.J., and B. Kaiser. 2003. Regulation of muscarinic acetylcholine receptor signaling. Pharmacology & therapeutics 98:197–220.

Waldhoer, M., S.E. Bartlett, and J.L. Whistler. 2004. OPIOID RECEPTORS. Annual Review of Biochemistry 73:953–990.

Wang, X., C. Wang, J. Zeng, X. Xu, P.Y.K. Hwang, W.-C. Yee, Y.-K. Ng, and S. Wang. 2005. Gene Transfer to Dorsal Root Ganglia by Intrathecal Injection: Effects on Regeneration of Peripheral Nerves. Mol Ther 12:314–320.

Wang, Y., D.M. Lonard, Y. Yu, D.C. Chow, T.G. Palzkill, J. Wang, R. Qi, A.J. Matzuk, X. Song, F. Madoux, P. Hodder, P. Chase, P.R. Griffin, S. Zhou, L. Liao, J. Xu, and B.W. O’Malley. 2014. Bufalin is a potent small-molecule inhibitor of the steroid receptor coactivators SRC-3 and SRC-1. Cancer research 74:1506–1517.

Wehrens, X.H., and A.R. Marks. 2004. Novel therapeutic approaches for heart failure by normalizing calcium cycling. Nat Rev Drug Discov 3:565–573.

Whistler, J.L., and M. von Zastrow. 1998. Morphine-activated opioid receptors elude desensitization by β-arrestin. Proceedings of the National Academy of Sciences of the United States of America 95:9914–9919.

Williams, J.T., S.L. Ingram, G. Henderson, C. Chavkin, M. von Zastrow, S. Schulz, T. Koch, C.J. Evans, and M.J. Christie. 2013. Regulation of µ-Opioid Receptors: Desensitization, Phosphorylation, Internalization, and Tolerance. Pharmacological Reviews 65:223–254.

Wu, W.L., Y.W. Lin, M.Y. Min, and C.C. Chen. 2010. Mice lacking Asic3 show reduced anxiety-like behavior on the elevated plus maze and reduced aggression. Genes, Brain and Behavior 9:603–614.

Ye, J., S. Chen, and T. Maniatis. 2011. Cardiac glycosides are potent inhibitors of interferon-β gene expression. Nature chemical biology 7:25–33.

Zaki, P.A., D.E. Keith, G.A. Brine, F.I. Carroll, and C.J. Evans. 2000. Ligand-Induced Changes in Surface µ-Opioid Receptor Number: Relationship to G Protein Activation? Journal of Pharmacology and Experimental Therapeutics 292:1127–1134.

Zeng, M.D.W., M.D.S. Dohi, M.D.H. Shimonaka, and M.D.T. Asano. 1999. Spinal Antinociceptive Action of Na+-K+Pump Inhibitor Ouabain and Its Interaction with Morphine and Lidocaine in Rats Anesthesiology 90:500–508.

Zheng, H., H.H. Loh, and P.-Y. Law. 2010. Agonist-selective signaling of G protein-coupled receptor: Mechanisms and implications. IUBMB Life 62:112–119.

